# Hand Knob Area of Motor Cortex in People with Tetraplegia Represents the Whole Body in a Modular Way

**DOI:** 10.1101/659839

**Authors:** Francis R. Willett, Darrel R. Deo, Donald T. Avansino, Paymon Rezaii, Leigh Hochberg, Jaimie Henderson, Krishna Shenoy

## Abstract

Decades after the motor homunculus was first proposed, it is still unknown how different body parts are intermixed and interrelated in human motor cortex at single-neuron resolution. Using microelectrode arrays, we studied how face, head, arm and leg movements on both sides of the body are represented in hand knob area of precentral gyrus in people with tetraplegia. Contrary to the traditional somatotopy, we found strong representation of all movements. Probing further, we found that ipsilateral and contralateral movements, and homologous arm and leg movements (e.g. wrist and ankle), had a correlated representation. Additionally, there were neural dimensions where the limb was represented independently of the movement. Together, these patterns formed a “modular” code that might facilitate skill transfer across limbs. We also investigated dual-effector movement, finding that more strongly represented effectors suppressed the activity of weaker effectors. Finally, we leveraged these results to improve discrete brain-computer interfaces by spreading targets across all limbs.

## Introduction

Neural activity in human motor cortex is traditionally thought to have a mixed somatotopic organization, in which face, arm and leg movements are represented in distinct areas of cortex but individual muscles within any one body area may be mixed (e.g. wrist and finger movements may overlap within arm area) (Schieber 2001). Classical stimulation studies in great apes and humans suggest an orderly arrangement of body parts along the precentral gyrus, with little intermixing between widely distinct areas of the body within any one individual (Leyton and Sherrington 1917; W. Penfield and Boldrey 1937; Wilder Penfield and Rasmussen 1950). fMRI studies also support the existence of an orderly map with largely separate face, arm and leg areas along the precentral gyrus and the anterior bank of the central sulcus (Lotze et al. 2000; Meier et al. 2008), as do electrocortiographic recordings in the high gamma band above precentral gyrus (albeit with occasional exceptions) (Crone, Miglioretti, Gordon, Sieracki, et al. 1998; Crone, Miglioretti, Gordon, and Lesser 1998; Miller et al. 2007; Ganguly et al. 2009; Ruescher et al. 2013). Nevertheless, the level of mixed representation within any one area of human precentral gyrus has not yet been quantified at single neuron resolution.

Here, we revisit motor somatotopy using microelectrode array recordings from arm/hand motor cortex (hand knob area (Yousry et al. 1997)) while two participants in the BrainGate2 pilot clinical trial made (or attempted) a variety of face, head, leg and arm movements. Surprisingly, we found strong and separable modulation for all tested movements. In one participant currently enrolled in the trial, we had a unique opportunity to probe further to characterize how movements of all four limbs are neurally represented in one area of human motor cortex. Previous fMRI and ECoG studies on human precentral gyrus have found that it contains a correlated representation of ipsilateral and contralateral arm/hand movements (that is, matching movements of the ipsilateral and contralateral arms have similar neural representations) (Diedrichsen, Wiestler, and Krakauer 2013; Wiestler, Waters-Metenier, and Diedrichsen 2014; Jin et al. 2016; Fujiwara et al. 2017; Bundy et al. 2018). Macaque studies also show a correlated representation of ipsilateral and contralateral reaching movements (Cisek, Crammond, and Kalaska 2003) that changes during bimanual movements (Rokni et al. 2003). Recently, a microelectrode array study of human parietal cortex found correlated representations of ipsilateral and contralateral arm movements (Zhang et al. 2017). Here, we examine these phenomena for the first time in human motor cortex with penetrating microelectrodes; we also extend these results to leg movements, revealing a neural code that links not only ipsilateral limbs to contralateral limbs, but also arm movements to homologous leg movements (e.g. wrist movements to ankle movements).

We call this neural code “modular” because it consists of separable movement-coding neural dimensions and limb-coding dimensions. In movement-coding neural dimensions, we found that movements were represented in a correlated manner across all four limbs. In limb-coding dimensions, the limb itself was represented largely independently of the movement. Together, these dimensions are indicative of a modular representation of movement that might underlie our ability to transfer motor skills between different limbs [as shown in (Kelso and Zanone 2002; Christou and Rodriguez 2008; Morris et al. 2009; Shea, Kovacs, and Panzer 2011)]; a motor skill could be learned in movement-coding dimensions and then transferred to another limb by changing activity in limb-coding dimensions.

In addition to examining single-effector movements, we also investigated how simultaneous movements of the arms, legs and head are represented. We find that neural activity nonlinearly changes during dual-effector movement, with the more strongly represented (primary) effector suppressing the activity of the weaker (secondary) effector. These neural changes might help to keep the secondary effector from interfering with the primary effector during simultaneous and independent control. Finally, we demonstrate, for the first time, a discrete intracortical brain-computer interface (BCI) that can decode movements across all four limbs. We show that this “full body” BCI improves information throughput relative to a single-effector approach.

## Results

### Tuning to Face, Head, Arm and Leg Movements

We used microelectrode array recordings from participants T5 and T7 to assess tuning to face, head, arm and leg movements in hand knob area of precentral gyrus. Participant T5 had a C4 spinal cord injury and was paralyzed from the neck down; he could move his face and head, but attempted arm and leg movements resulted in little or no overt motion. Participant T7 had ALS and could move all joints tested, although some of his arm movements were limited due to weakness.

In this experiment, T5 and T7 made (or attempted to make) movements in sync with visual cues displayed on a computer screen (Fig. 1A). T5 completed an instructed delay version of the task where each trial randomly cued one of 32 possible movements spanning the face, head, arm and legs. For face and head movements, T5 was instructed to move normally; for arm and leg movements, T5 was instructed to attempt to make the movement as if he were not paralyzed. T7, whose more restricted data was collected earlier in a different study (and who is no longer enrolled), completed an alternating paired movement task with a block design. Each block tested a different movement pair, during which T7 alternated between making each of the paired movements every 3 seconds.

**Figure 1.**
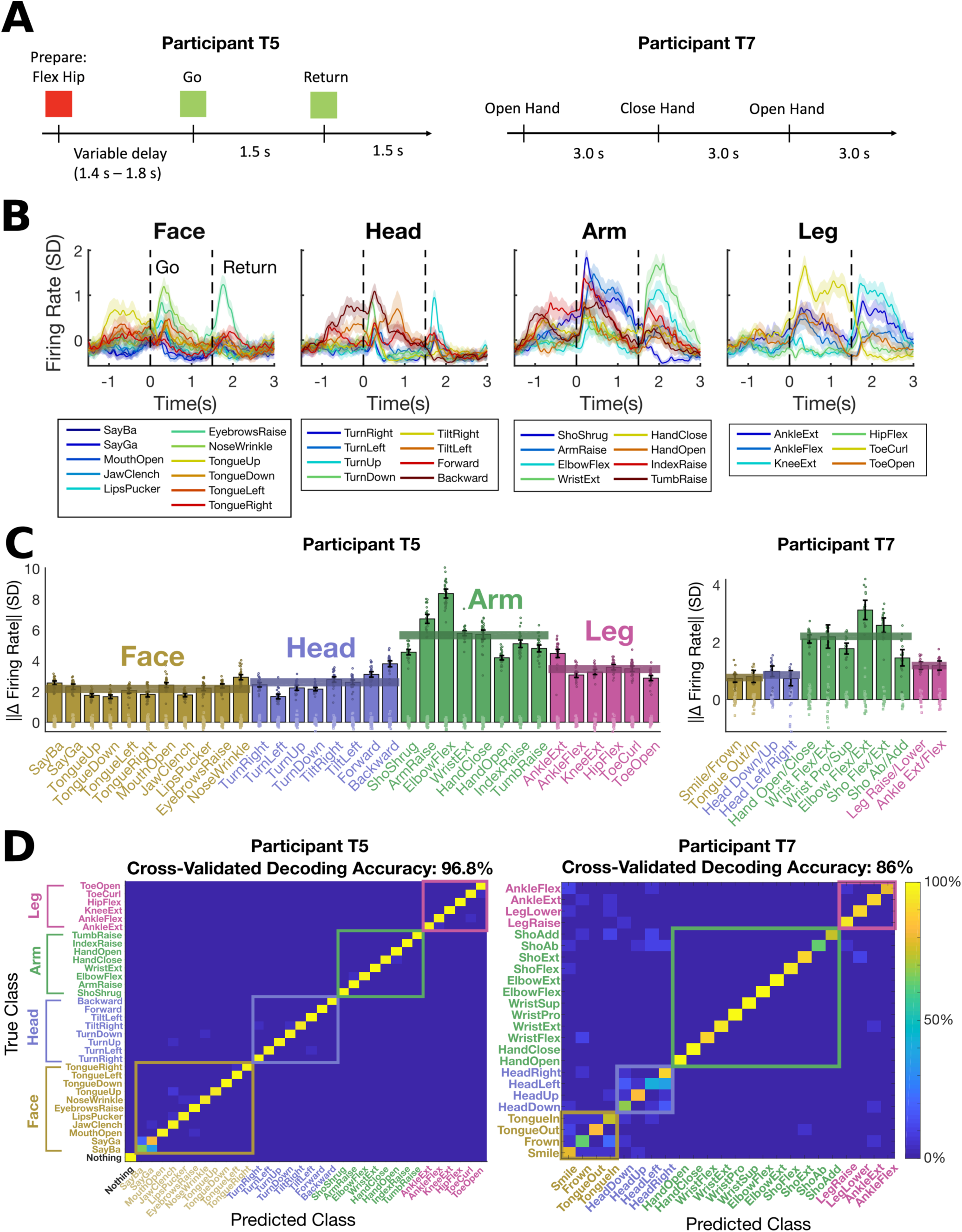
Separable and robust modulation for face, head, arm and leg movements. (A) Neural activity was recorded while participants T5 and T7 completed cued movement tasks that instructed them to make (or attempt to make) movements of the face, head arm and leg in sync with text appearing on a computer monitor. (B) The mean firing rate recorded for each cued movement is shown for an example electrode from participant T5. Shaded regions indicate 95% CIs. Neural activity was convolved with a Gaussian smoothing kernel (30 ms std). (C) The size of the neural modulation for each movement was quantified by comparing to a baseline “do nothing” condition (in participant T5) or by comparing to its paired movement (participant T7). Each bar indicates the size (Euclidean norm) of the mean change in firing rate across the neural population (with a 95% CI) and each dot corresponds to the cross-validated projection of a single trial’s firing rate vector onto the “difference” axis. Statistically significant modulation was observed for all movements (no confidence intervals contain 0). Translucent horizontal bars indicate the mean bar height for each movement type. (D) A Gaussian naive Bayes classifier was used to classify each trial’s movement using the firing rate in a window from 200 to 600 ms (T5) or 200 to 1600 ms (T7) after the go cue. Confusion matrices are shown from offline cross-validated performance results; each entry (i, j) in the matrix is colored according to the percentage of trials where movement j was decoded (out of all trials where movement i was cued).

Despite recording from microelectrode arrays placed in the hand knob region of precentral gyrus, we found strong neural tuning to all tested movements, including those of the face, head and leg. Fig. 1B shows an example electrode from T5 that was tuned to all movement categories. Here, and in most other results, we analyzed binned “threshold crossing” firing rates for each microelectrode channel (although see Supplemental Figure 3, mentioned below, which reports results from spike-sorted single units). Analyzing threshold crossings as opposed to spike-sorted units allowed us to leverage information from more electrodes, since many electrodes recorded activity from multiple neurons that could not be precisely distinguished from each other. Recent results indicate that neural population structure can be accurately estimated using threshold crossing rates alone (Trautmann et al. 2017, 2019). Binned threshold crossing rates were z-scored by subtracting the mean and dividing by the standard deviation (and are reported in units of SD).

In Fig. 1C we summarize the modulation observed for each individual movement (T5) or pair of movements (T7) across all electrodes from both microelectrode arrays. For participant T5, we quantified modulation size by computing the magnitude (Euclidean norm) of the difference between the mean firing rates observed during a “do nothing” control condition and the mean rates observed during the movement of interest. Modulation size was estimated in a cross-validated, unbiased way (see Methods). For participant T7, we computed the magnitude of the difference between the mean firing rates observed for each pair of movements. Firing rates were computed for each trial in a time window 200 to 600 ms after the go cue (T5) or 200 to 1600 ms after the go cue (T7). Fig. 1C shows robust modulation for each tested movement in both participants (the clusters of single trial firing rates are clearly separable and the confidence intervals are far from zero). Tuning to arm movement was the strongest in both participants, with tuning to non-arm movements 38% (face), 46% (head) and 61% (leg) as large as arm movement tuning in T5, and 38% (face), 34% (head) and 53% (leg) as large in T7.

Next, we tested whether the modulation for each movement was distinguishable from other movements by using a cross-validated naive Bayes classifier to decode the movement on each trial. The high classification accuracies (Fig. 1D) confirm that the tuning to each movement was unique and separable in both participants (as opposed to being caused by a non-specific signal that was the same for all movements). In Supplemental Figure 1, we plot the neural activity in the top neural dimensions found using principal components analysis (PCA) to illustrate that, in both participants, the activity is rich and multi-dimensional for all movement types.

Finally, we searched for somatotopy across the microelectrode arrays and saw no clear patterns; tuning to all four movement types was highly intermixed in both participants and many electrodes were tuned to multiple movement types (Supplemental Figure 2). We also analyzed well-isolated, spike-sorted single units and found that they were frequently tuned to multiple types of movement as well (Supplemental Figure 3).

### A Modular Neural Code for Arm and Leg Movements

Next, we assessed the tuning to ipsilateral attempted arm and leg movements in participant T5 (Fig. 2A). Our remaining results (Figs. 2-6) are based on data from T5, who is currently enrolled in the trial. Fig 2A shows that firing rates changed two-thirds as much for ipsilateral movement as compared to contralateral movement (65% as much for the arm, 78% as much for the leg). Tuning to ipsilateral movements was separable and robust; a cross-validated naive Bayes classifier could decode the movement type with 97.4% accuracy (from amongst all those shown in Fig. 2A) using a 200 to 600 ms window after the go cue. In addition to the *magnitude* of firing rate change, we also assessed the *signed* change in firing rate across all electrodes (Fig. 2B). For both the arms and the legs, contralateral attempted movements evoked a larger global increase in firing rate than did ipsilateral attempted movements.

**Figure 2.**
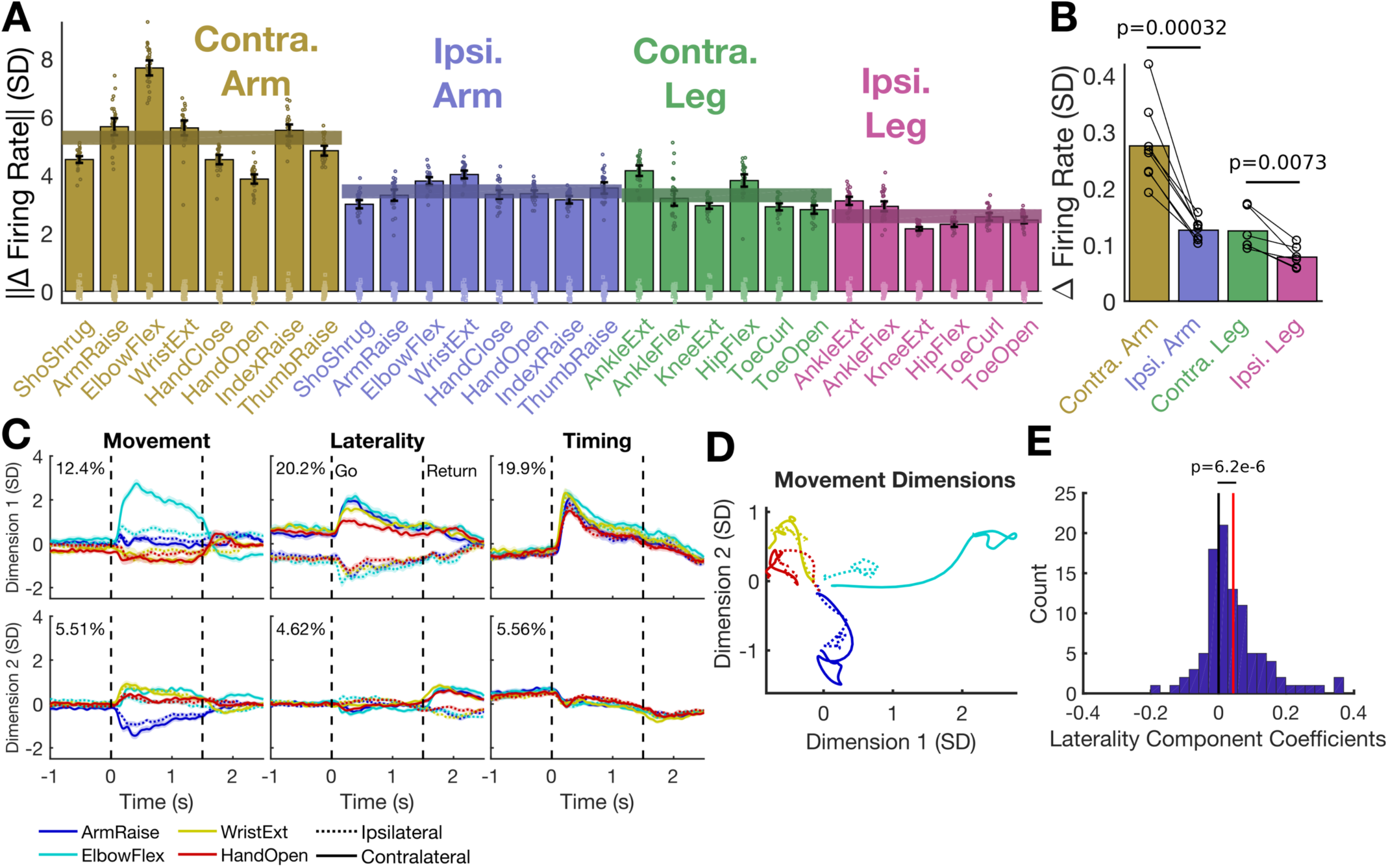
Representation of attempted ipsilateral and contralateral arm and leg movements. (A) The size of the neural modulation for each attempted movement was quantified by comparing to a baseline “do nothing” condition. Each bar indicates the size (Euclidean norm) of the mean change in neural population activity (with a 95% CI) and each dot corresponds to the cross-validated projection of a single trial’s neural activity onto the “difference” axis. Translucent horizontal bars indicate the mean bar height for each attempted movement type. (B) The average *signed* change in firing rate per electrode is shown for each movement (black circle) and each effector (bar). Averages across all movements are indicated by bar height. Lines indicate pairwise comparisons between matching ipsilateral and contralateral attempted movements. Change in firing rate was measured with respect to a “do nothing” control condition. P-values were computed with paired t-tests. (C) To reveal how the attempted movements in A were neurally represented in relation to each other, demixed PCA was applied to a set of four example attempted arm movements. Each panel shows the mean activity for a single dimension and each column shows dimensions related to a particular factor (marginalization). The movement and laterality interaction was included in the movement factor. Each trace shows the mean activity for a single condition and the shaded regions indicate 95% CIs. The percent variance explained by each dimension is indicated in the top left. (D) The mean activity through time in the first two movement dimensions is plotted as a trajectory for each condition; the dashed lines lie mostly near the solid lines and travel in the same direction, indicating that activity for a given movement is similar across sides of the body. (E) The distribution of each electrode’s coefficient in the laterality component is positive on average (red line indicates the mean and black line indicates zero), meaning that this dimension describes global increases and decreases in firing rate (e.g. those shown in B).

We then sought to probe the structure of the neural tuning we observed. We were guided by prior single unit recordings in macaques (Cisek, Crammond, and Kalaska 2003; Rokni et al. 2003) and fMRI and ECoG studies in people (Diedrichsen, Wiestler, and Krakauer 2013; Jin et al. 2016; Bundy et al. 2018) which found that ipsilateral and contralateral movements had correlated representations in motor cortex. If this is the case, we hypothesized that there should also exist laterality-related neural dimensions which code for the side of the body independently of the movement details. These laterality dimensions would help downstream areas to distinguish between contralateral and ipsilateral movement. This kind of “modular” neural code, as defined by correlated movement-coding dimensions and independent effector-coding dimensions, is distinct from a muscle-like representation. In a muscle-like representation, the neural representation of contralateral and ipsilateral movements would exist in largely orthogonal dimensions and hence be uncorrelated (as recently found in macaque M1 (Ames and Churchland 2019; Heming et al. 2019)), since the muscles are largely non-overlapping.

We used demixed PCA (dPCA)(Kobak et al. 2016) to visualize the neural activity in a way that shows where the neural activity falls on the spectrum between “muscle-like” and “modular”. dPCA decomposes the neural data into a set of dimensions that each explain variance related to one particular marginalization of the data. We marginalized the data according to three separate factors: time, laterality, and movement. In the movement marginalization, we included the main effect of the movement condition as well as the interaction between movement and laterality. If the muscle-like hypothesis holds, there should be small laterality dimensions and large movement dimensions in which the activity is dissimilar for the same movements on opposite sides of the body. On the other hand, in the case of modular coding, there should be a large laterality dimension (in addition to large movement dimensions), and activity in the movement dimensions should be similar for the same movements on opposite sides of the body.

Fig. 2C-E shows the result of the dPCA analysis. The data appear more consistent with the modular coding hypothesis. There is a large laterality dimension representing the side of the body where the movement occurred independently of the movement itself (Fig. 2C, middle column). Additionally, activity in the movement dimensions (Fig. 2C, left column) is similar for any given movement (e.g. wrist extension) regardless of which side of the body it is on (Fig. 2D). Finally, we examined how individual electrodes mapped onto the laterality dimension and found that they had positive coefficients on average (Fig. 2E). That is, activity in the laterality dimension corresponded to a global increase in firing rate for contralateral movements, accounting for Fig. 2B.

Next, we tested for modularity more comprehensively. First, we used a cross-validation approach to confirm that the large laterality dimension revealed in Fig. 2C was consistent for held-out movements that were not used to find it. Fig. 3A shows that activity in the laterality dimension correctly separates attempted movements made on opposite sides of the body even for held-out single movements (diagonal panels) and for held out effectors (i.e. arm vs. leg, off-diagonal panels). We also used cross-validated dPCA to search for a putative “arm vs. leg” dimension that codes for whether a movement was attempted with the arm or the leg (independently of the movement details or the side of the body on which it occurs). Results indicate the presence of an arm vs. leg dimension in addition to a laterality dimension. Both the laterality and arm vs. leg dimensions contributed to an overall increase in signed firing rate for contralateral (vs. ipsilateral) and arm (vs. leg) movements; nevertheless, these neural dimensions were clearly separable from each other and from neural dimensions related to timing (Supplemental Figure 4).

**Figure 3.**
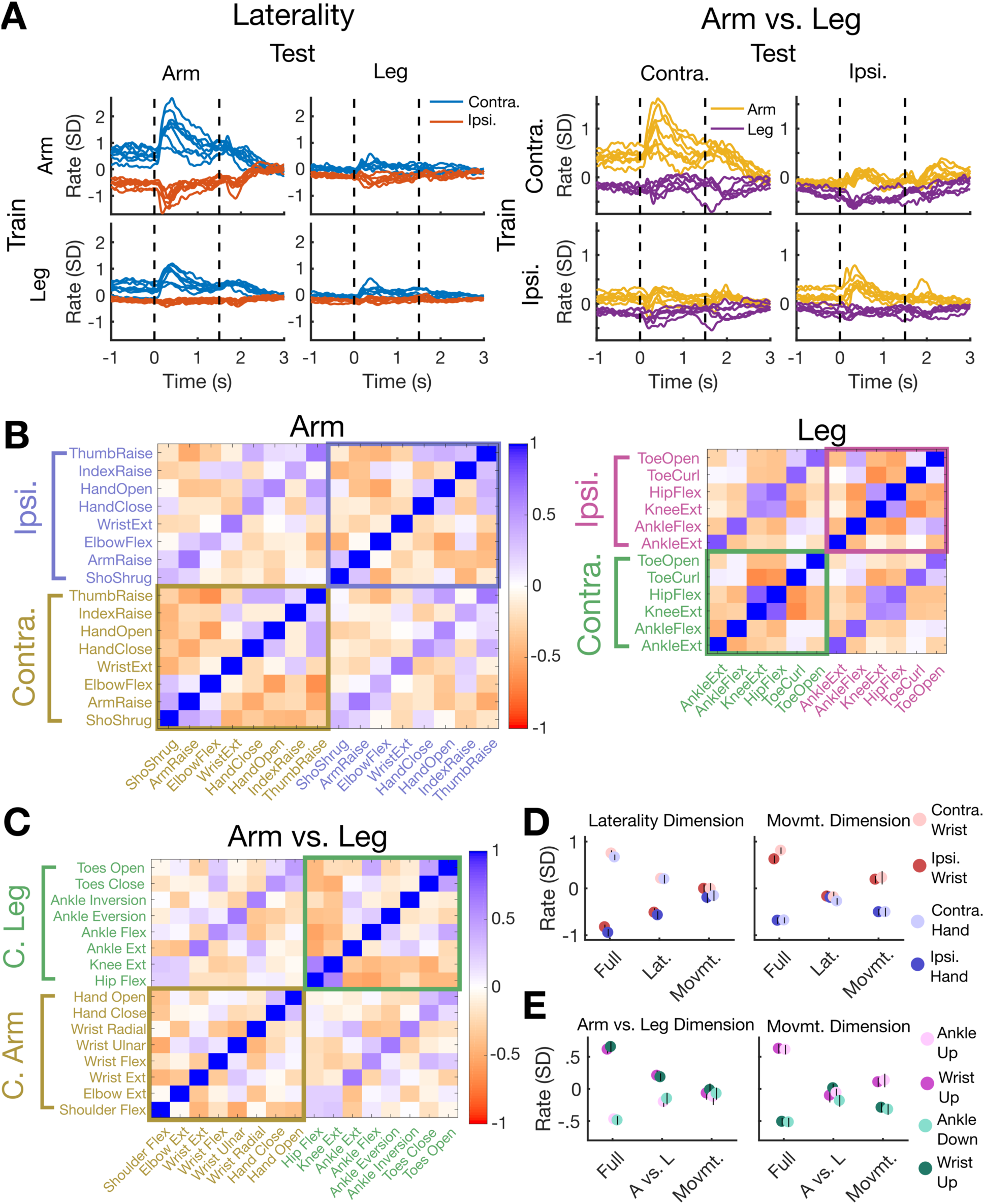
A modular code for attempted arm and leg movements. (A) A cross-validation analysis demonstrates the existence of neural dimensions that code for laterality (left vs. right side of the body) and effector (arm vs. leg). Each dimension was identified using only a subset of the data (rows) and then applied to all data (columns). In diagonal panels (e.g. train on arm and test on arm), each individual movement was held out (that is, each movement’s trace was made using dimensions found using other movements only). (B) Each square *(i,j)* of the correlation matrices indicates the correlation coefficient observed when correlating the mean firing rate vector of movement *i* with that of movement *j*. The off-diagonal banding shows that ipsilateral and contralateral movements were correlated with each other, for both the arm (left) and leg (right). (C) Correlations between pairs of contralateral arm and leg movements were computed to reveal a surprising pattern: homologous movements of the arm and leg are correlated (e.g. hand opening and closing is uniquely correlated with toe opening and closing, as indicated by the off-diagonal band). This data is from a different dataset than that used in B, and was designed specifically to test many pairings of homologous arm and leg movements. (D) Neural activity in laterality-coding and movement-coding dimensions is partially active during an instructed delay period where only the laterality or the movement is specified. Each panel corresponds to a different neural dimension. Each circle indicates the mean (and its black line indicates the 95% CI) of activity in each dimension in the case where the full movement is specified (Full), only the laterality is specified (Lat.), or only the movement is specified (Movmt.). A 300 ms time window preceding the go cue was used for analysis. (E) Same as in D, showing results from a different experiment testing the partial preparation of arm vs. leg (wrist vs. ankle) and movement direction.

Next, we quantified the correlations between firing rate vectors observed for attempted arm and leg movements on opposite sides of the body (Fig. 3B). First, firing rates were averaged across trials in a window between 200 and 600 ms after the go cue and then concatenated across electrodes into a vector. Then, the mean firing rate vector across all movements within each side of the body was subtracted to remove the effect of the laterality dimension (otherwise, all movements on different sides of the body would appear negatively correlated). Finally, the correlation (Pearson’s r) between the firing rate vectors for each pair of movements was computed. We observed a consistently positive correlation between matching ipsilateral and contralateral movements for both the arms and legs (indicated by the off-diagonal banding in Fig. 3B). The correlation values for matching movements, such as contralateral vs. ipsilateral wrist extension, were statistically significantly greater than those for non-matching movements, such as contralateral wrist extension vs. ipsilateral elbow flexion (mean r=0.52 vs. r=-0.07 for the arm, p=1.1e-10; r=0.73 vs. r=-0.14 for the leg, p=2.3e-09; significance assessed with a t-test). All correlations were computed in a cross-validated, unbiased manner (see Methods).

Surprisingly, attempted homologous movements of the arm and leg were also more correlated than non-homologous movements (e.g. hand grasp and toe curl are positively correlated, while hand close and ankle inversion are not, as shown in the off-diagonal band in Fig. 3C). Correlations between homologous movements were statistically significantly greater than for non-homologous movements (mean r=0.48 vs. r=-0.06, p=2e-10; significance assessed with a t-test). We were also able to confirm this result for a more limited set of four arm and leg movements in participant T7 (mean r=0.22 vs. r=-0.07, p=0.0036, Supplemental Figure 5). These homologous correlations, along with the arm vs. leg dimension shown in Fig. 3A, extend the modular neural coding scheme to pairs of arm and leg movements.

Finally, we examined modular coding through the lens of preparatory activity. We reasoned that if modular coding was the result of independent specification of the movement type and the limb to be moved, then neural activity in limb-coding dimensions should be able to be prepared independently of neural activity in movement-coding dimensions in the case of partial information (i.e. when only the limb or only the movement is cued in an instructed delay task). We tested this by having T5 perform a Partial Cue Task (see Methods). We found that when only the movement type is specified during the instructed delay period (i.e. “wrist extension” or “hand close”), activity in the movement dimension partially codes for the upcoming movement while activity in the laterality dimension doesn’t change (Fig. 3D, top right panel). Likewise, when only the laterality is specified (i.e. “left” or “right”), activity in the laterality dimension codes for the upcoming side of the body while activity in the movement dimension doesn’t change (Fig. 3D, top left panel). This result also holds for partial specification of arm vs. leg and movement direction (Fig. 3E). By examining the single trial distributions of preparatory activity (Supplemental Figure 6), we confirmed that the neural activity shows patterns consistent with genuine partial preparation as opposed to a mixture of full preparation and no preparation.

### Arms and Legs Share an Intrinsic Representation of Direction

Figures 1-3 assess neural modulation largely to movements around a single joint. Here, we assess how movement direction is represented across different effectors. Participant T5 completed an instructed delay, spatially cued movement task which randomly cued one of eight directional movements from one of four effectors (Fig. 4A). We confirmed that neural activity was linearly tuned to movement direction [as expected from previous work (Georgopoulos et al. 1982)] by plotting the mean firing rates for each movement in the first two principal components (in a time window between 200 to 1000 ms after the go cue); cross-validation was used to ensure an unbiased result. The neural activity forms a ring which matches the geometry of the targets for each of the four body parts tested and is indicative of linear tuning to direction (Fig. 4B).

**Figure 4.**
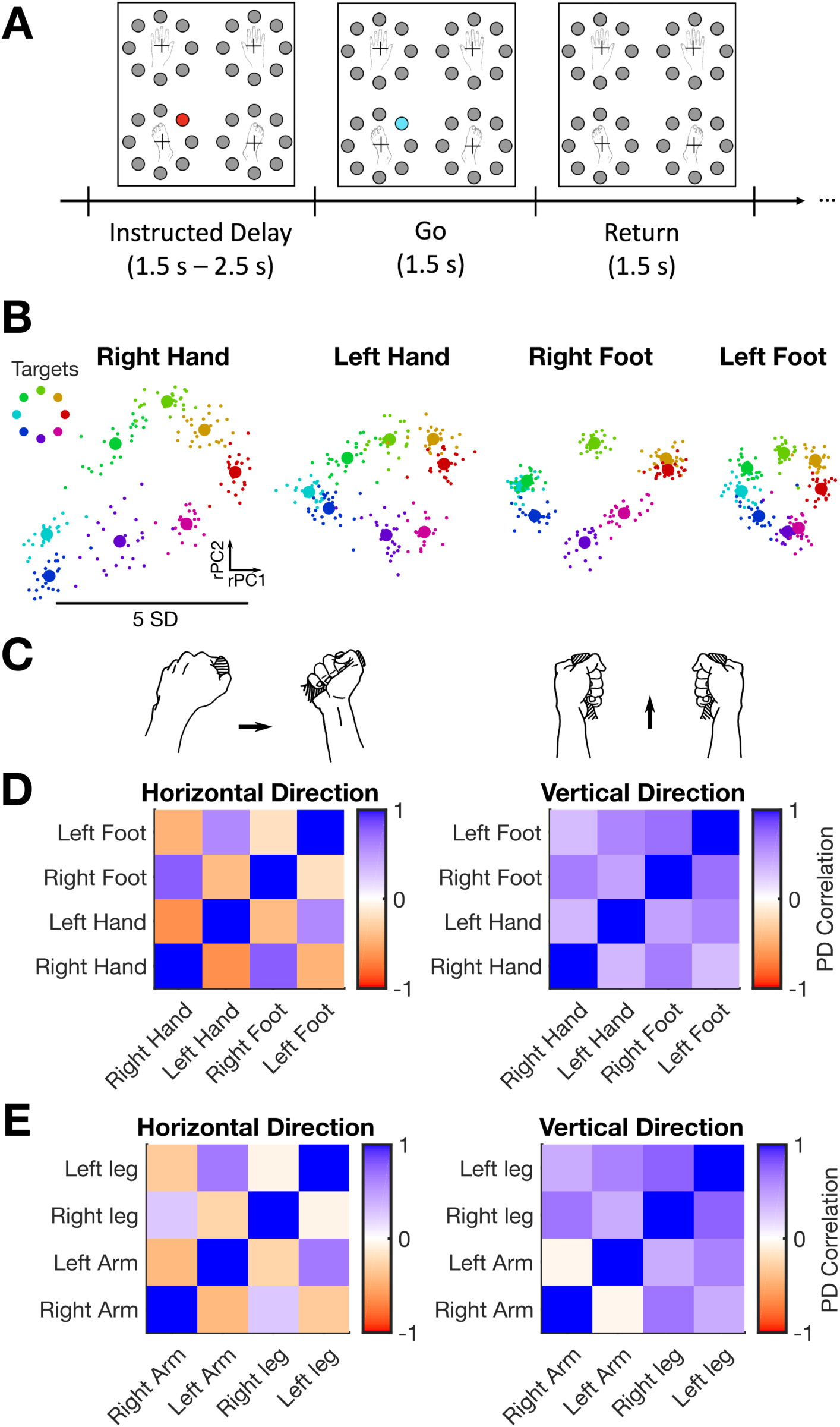
The representation of movement direction across the arms and legs is intrinsic. (A) An instructed delay, spatially cued movement task was used to probe the neural representation of directional movements across different effectors. The task illustrated here cues directional movements of the hand (as if pointing a joystick) and foot (pointing the ankles) on both sides of the body. (B) Mean (large circles) and single trial (small circles) averaged firing rates during the “go” period of a radial 8 directional movement task are plotted in the first two dimensions found by PCA. The dimensions are rotated to align with the targets and thus are labeled as “rPC1” and “rPC2”. (C) Illustration of how horizontal movements in the same extrinsic direction (left image) require opposite joint movements while vertical movements in the same direction (right image) require identical joint movements. Hands are depicted making joystick pointing movements (matching T5’s instructions). (D,E) Directional tuning was correlated across effectors in the horizontal and vertical directions. Each smaller square (i,j) indicates the correlation coefficient between the horizontal or vertical tuning coefficients that were fit to each body part i and j separately using a linear model. In the vertical direction, the directional tuning across all effectors is positively correlated. In the horizontal direction, however, effectors on opposite sides of the body are negatively correlated, indicating that movement direction is intrinsically represented. Movement in the horizontal direction on opposite sides of the body requires opposite joint movements to move in the same extrinsic direction.

Next, to assess how neural tuning to movement direction was correlated across different body parts, we first fit a linear tuning model for each body part separately to obtain horizontal and vertical tuning coefficients (preferred directions) for each electrode. Then, we correlated these horizontal and vertical coefficients separately between each different body part to see if a representation of movement direction was shared across body parts (Fig. 4C-D). If an “extrinsic” (visuospatial) representation were shared, then both the horizontal and vertical coefficients should be positively correlated. On the other hand, if an “intrinsic” (joint or muscle-related) representation were shared, then the vertical coefficients should be positively correlated in all cases (since these correspond to the same joint movements), while the horizontal coefficients should be negatively correlated across different sides of the body (since these correspond to opposite joint movements) but positively correlated on the same side of the body. We found that, across both the wrists and ankles (Fig. 4C) and the arms and legs (Fig. 4D), the pattern of correlations was more consistent with a shared intrinsic representation (mean r=0.50 for cases of matching joint movements, mean r=-0.34 for cases of opposite joint movements, p=1.4e-17; significance assessed with a t-test).

### Changes in Neural Tuning during Simultaneous Movement of Two Effectors

We hypothesized that the modular code we observed for single-effector movements, consisting of strong, correlated tuning for all four limbs in a single area, might not be ideal for controlling multiple effectors at the same time (since commands for one limb might be interpreted as commands for the other). Prior work on bimanual reaching in macaques showed a decorrelation and suppression of ipsilateral-related neural activity during bimanual reaches (Rokni et al. 2003). To investigate this possibility, we tested all pairings of the following five effectors: left arm, right arm, left leg, right leg, and head. For each effector pair, we used an instructed delay cued movement task (Fig. 5A) to probe the neural representation of left and right directional movements made either in isolation or simultaneously. For all pairs, we observed separable neural modulation for each of the four simultaneous movement conditions; a cross-validated naive Bayes classifier was able to distinguish between these conditions with an accuracy ranging from 86.1% to 100% (mean=95% across all pairs) using the firing rates within a 200 to 1000 ms window after the go cue.

**Figure 5.**
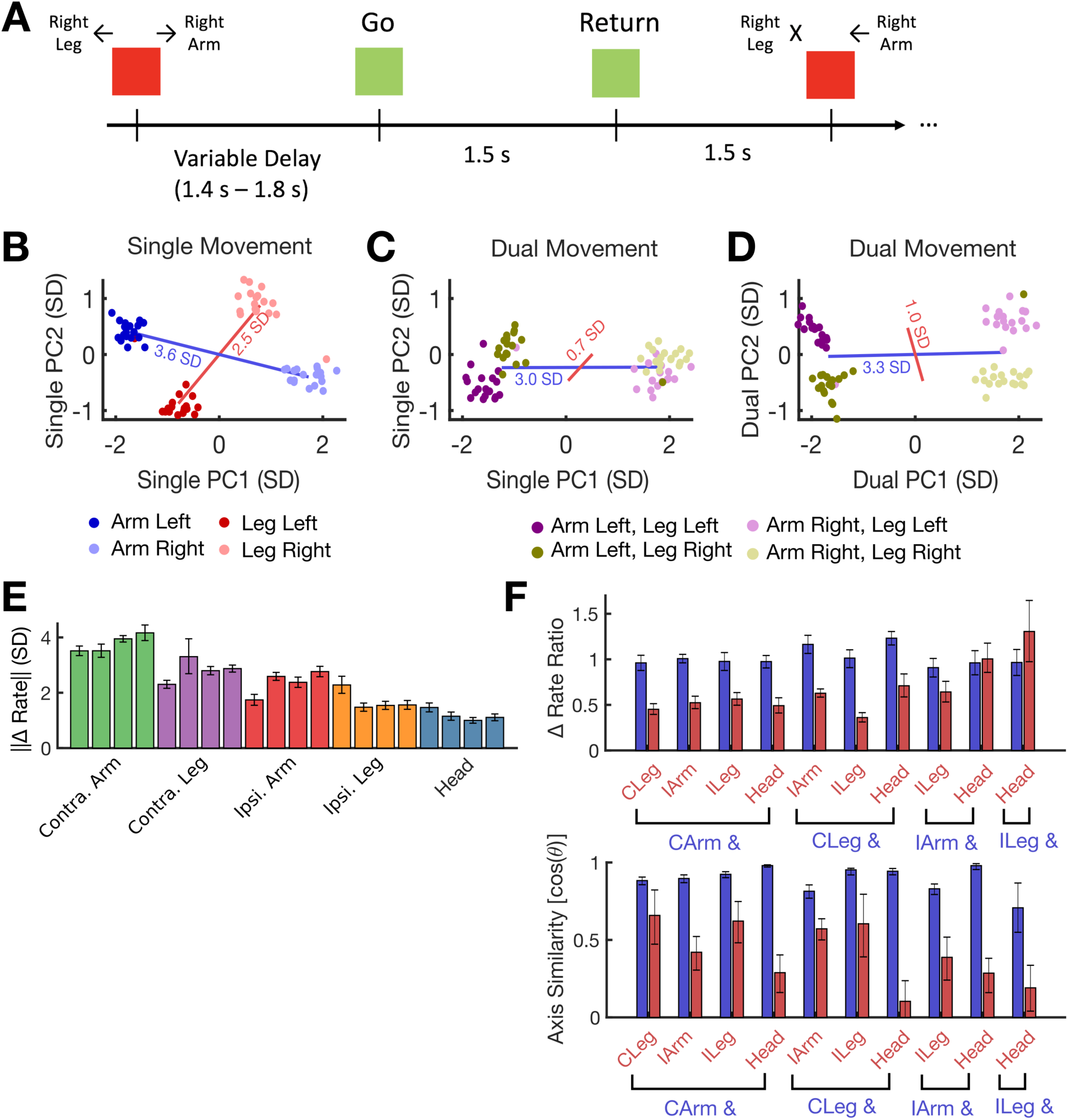
Representation of the “secondary” effector changes and becomes weaker during dual movement. (A) An instructed delay movement cue task was used to probe the neural representation of dual (multi-effector) movement. In this example experiment, participant T5 attempted to move his contralateral arm and leg to the left or right either simultaneously (e.g. the first illustrated movement) or individually (e.g. the second illustrated movement). (B) Example single trial firing rates vectors during the “go” period are plotted in the top two neural dimensions that show differences in firing rate due to movement direction. Red and blue lines connect their respective cluster means and their lengths are a measure of tuning strength. (C) The firing rates observed during dual movement are plotted in the same dimensions used in B. Modulation for the leg is considerably weakened in this space (the red line is smaller than in B). Red and blue lines show the distance between the dark and light clusters (blue, arm movement) and the purple and gold clusters (red, leg movement). (D) The data in C is plotted in the top two principal components that describe the dual movement data. The modulation for the leg appears larger, indicating that the axis on which it is represented has changed relative to single movement. (E) Each bar depicts, for a single experiment, the mean change (± 95% CI) of firing rate observed during single movement for each effector (e.g. the blue and red lines in B). (F) The ratio of dual/single modulation (top) and the change in axis during dual movement (bottom) is shown for each individual experiment. Each pair of bars is from a separate experiment comparing two different body parts. The blue bar corresponds to whichever body part of the two tested that had the largest modulation in E. Results show that the “secondary” effector (red bar) almost always decreased its modulation more sharply during simultaneous movement (top) and changed its axis of representation more (bottom).

Fig. 5B shows example data from one effector pairing (contralateral arm and leg). To visualize the neural activity, firing rate vectors were computed for each trial using a 200 ms to 1000 ms time window after the go cue. Then, two neural dimensions were found in which to visualize these vectors by using PCA to highlight differences in firing rate due to movement direction (see Methods for details). The example data shows clear and separable firing rate clusters for each movement when the effectors are moved in isolation.

Next, we used these same two dimensions to visualize the neural activity observed during dual effector movement (Fig. 5C). Interestingly, the neural representation of arm movement appears largely intact (the blue line is similar) while the representation of leg movement shrinks considerably in this space (the red line is much smaller). This could be due to the fact that the axis of representation has changed and no longer aligns with the single movement PCs. In Fig. 5D we show the same activity but in neural dimensions that best explain dual movement (found again using PCA). Although the modulation for leg movement is still smaller than in Fig. 5B, it is enlarged relative to Fig. 5C, indicating that both the size of the modulation has decreased and the axis of representation has shifted.

We then summarized the effects observed in Fig. 5B-D across all effector pairings (Fig. 5E-F). Fig. 5E summarizes the magnitude of the change in firing rate caused by single effector movement when each effector is moved separately (e.g. the red and blue lines in Fig. 5B). Contralateral arm movements cause the largest change in firing rate, followed by the contralateral leg, ipsilateral arm, ipsilateral leg and head. Fig. 5F shows, for each effector pairing, how the size and the axis of the neural representation changed for each effector during simultaneous movement. To measure how the size changed, we divided the size of modulation during dual movement (the length of the red and blue lines in Fig. 5C) by the size of modulation during single movement (the length of the red and blue lines in Fig. 5B). Values less than 1 indicate that the neural activity was attenuated. A pattern holds across most pairings whereby the neural representation of the “primary” effector (blue bars) stays relatively constant while the representation of the “secondary” effector (red bars) is attenuated.

Fig. 5E also shows the same pattern for the change in axis; the primary effector mostly retains its axis of representation while the representation of the secondary effector changes more. The change in axis was measured by computing the cosine of the angle between the single movement axis and the dual movement axis. We found that this change in axis caused, on average, the axes of representation for the two effectors to become less correlated with each other (Supplemental Figure 7). The suppression and decorrelation effects observed here might help to facilitate independent dual-effector control by working to keep the somatotopically dominant effector’s activity free from interference.

### Leveraging Multiple Effectors Increases the Performance of a Discrete Decoding BCI

Current intracortical brain-computer interfaces (iBCIs) are limited to recording from a small number of locations on the cortex [e.g. (Hochberg et al. 2012; Collinger et al. 2013; Aflalo et al. 2015; Bouton et al. 2016; Pandarinath et al. 2017; Ajiboye et al. 2017)], making it difficult to record from both the arm and leg areas of precentral gyrus across both hemispheres. The results above, which show a strong representation of all four limbs in only a small patch of cortex, opens up the possibility of decoding movements from all four limbs to improve iBCI performance. One context in which this activity could be useful is to enable accurate selection from a large number of targets or keys by mapping distinct movements across all four limbs to different targets. This approach might improve performance relative to system that only uses the contralateral arm, since neural activity patterns from other limbs might be more distinct from each other.

We tested this idea with a discrete decoding iBCI that decoded, in closed-loop, which movement out of a set of N movements was selected by the user. We focused on directional movements made by the wrists (as if pointing a joystick) and the ankles (pointing the foot), consistent with prior work on discrete iBCIs that also decoded directional movements (Musallam et al. 2004; Santhanam et al. 2006). Fig. 6A shows the three target configurations tested. In all cases, the tasks were performed in a “time-locked” fashion where each trial lasted one second. For each trial, a target would illuminate and T5 had exactly one second to identify the target and perform the indicated movement which was then decoded at the end of the one second mark. After decoding, a sound played indicating to T5 whether the target was correctly decoded or not and the next target immediately appeared.

First, we motivate the need for using multiple effectors by showing how information throughput [measured with “achieved bit rate” (Nuyujukian et al. 2015)] and decoding accuracy varies as a function of the number of targets (Fig. 6B). After only 6 targets, accuracy and bit rate start to decrease when using a single contralateral effector. Next, we tested offline whether spreading out 16 targets across multiple effectors could increase performance (Fig. 6 C-E). When spread across all four limbs (4 targets per limb), accuracy is near 100% (Fig. 6C). The improvement in accuracy is a result of increasing the distance between each target in neural population state space (Fig. 6D) and the dimensionality of the neural population space spanned by the targets (Fig. 6E). We then confirmed online that information throughput was higher when using multiple effectors. For this experiment, we first performed offline simulations to select the optimal number of targets for the single effector, dual effector, and quad effector configurations. Even when using an optimized number of targets, the four-effector layout led to improved performance on average (Supplemental Video 1 shows example trials). Decoding errors mostly confused targets from the arm and leg on the same side of the body (Fig. 6G), presumably due to considerable correlations in directional modulation (as shown in Fig. 4).

**Figure 6.**
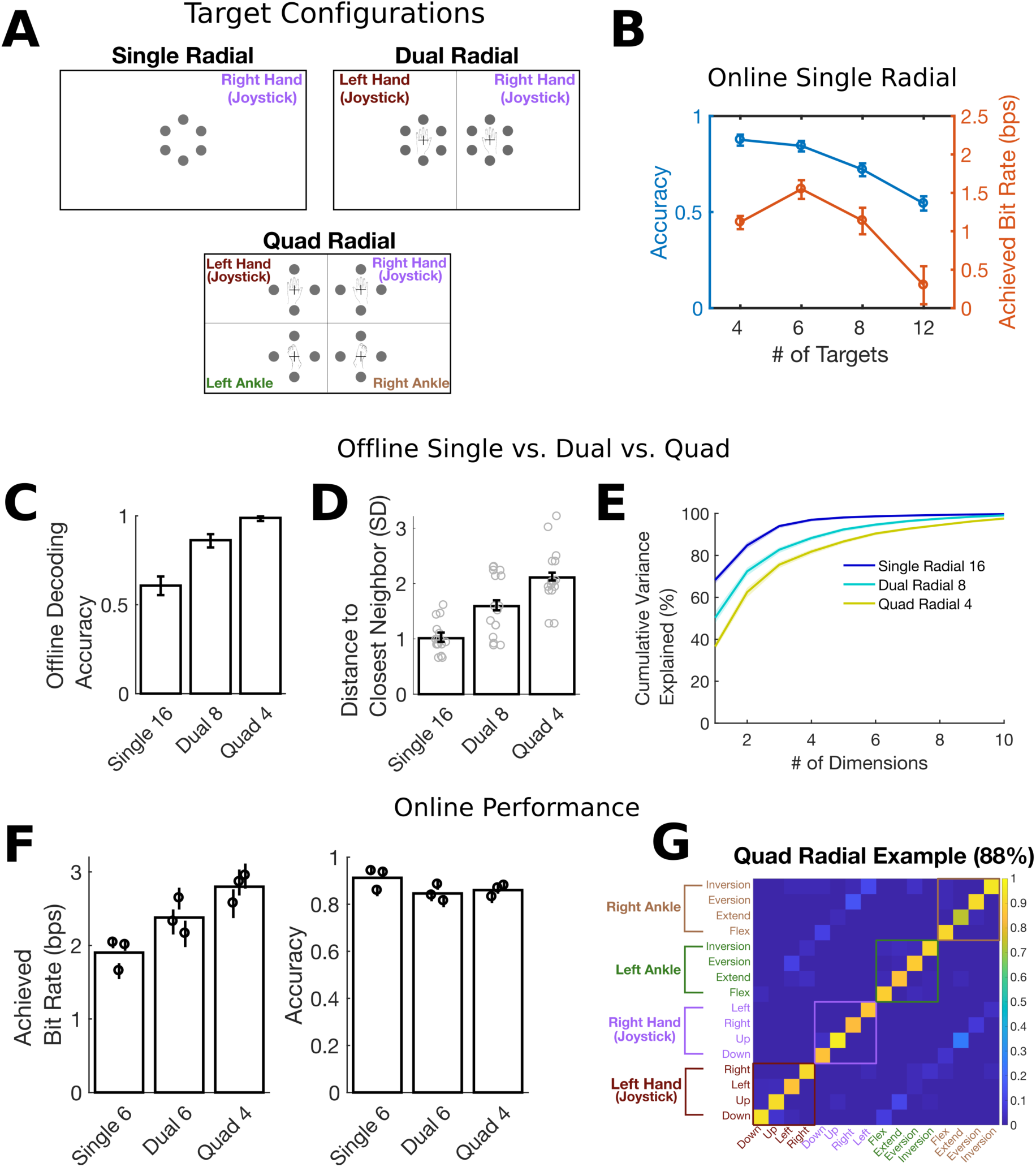
Spreading targets across multiple effectors increases the decoding accuracy and information throughput of a discrete decoding brain-computer interface. (A) Target configurations for the three discrete decoding tasks. Each target was associated with a specific attempted movement. (B) Online discrete decoding performance for a single contralateral effector (right wrist) as a function of the number of radially spaced targets included in the task. Bit rate and accuracy decline if more than 6 targets are used. (C) Offline decoding accuracy significantly increases as targets are spread across more effectors (mean and 95% CI shown). (D) As the number of effectors increases, for each target the distance to its closest neighbor in neural population state space increases (each gray dot corresponds to a single target and indicates the distance to its closest neighbor in neural population state space). The bar height indicates the mean of these dots and the error bar shows the 95% CI of the mean. (E) The dimensionality of the neural activity increases with the number of effectors. Cumulative variance explained is plotted (mean ± 95% CI) as a function of the number of PCA components used to explain the time-varying, mean neural activity across all targets. (F) Online discrete decoder performance metrics for target layouts shown in A. The achieved bitrate significantly increases with the number of effectors. Each circle shows the mean performance (± 95% CI) achieved during a single session; bar heights show the mean across all sessions. (G) Example confusion matrix from an online discrete decoding Quad Radial task. Each entry (i, j) in the matrix is colored according to the fraction of trials where movement j was decoded (out of all trials where movement i was cued). Off-diagonal banding shows that most errors were made when classifying between matching hand and foot movements.

Finally, we explored spreading targets across more body parts than just the wrists and ankles (including the elbow, knee, hip, toes, and fingers). Supplemental Figure 8 and Supplemental Video 2 show that high decoding accuracies (mean of 95%) can be achieved across 32 targets in this manner, as long as an instructed delay period is added to give T5 time to recognize the target and prepare the movement.

## Discussion

We found, in two participants, that there was strong neural representation of face, head, arm and leg movements in hand knob area of precentral gyrus (although modulation was strongest for attempted arm & hand movements, as expected). In one participant, we probed further to characterize, for the first time at microelectrode resolution, the structure of the neural code for movements across the whole body in one area of precentral gyrus. We found that arm and leg movements were represented in a correlated way. Movements of the same body part on different sides of the body, and homologous movements of the arm and leg (e.g. hand grasp and toe curl), were significantly correlated in neural dimensions that coded for movement type. These correlations appeared to be largely in an intrinsic (joint-based) frame as opposed to an extrinsic (visuospatial) frame. In other neural dimensions, the limb itself was represented largely independently of the movement; activity in these dimensions caused firing rates to be lower overall for ipsilateral movements and leg movements. Together, the activity in these neural dimensions form a partially “modular” neural code for movement that differs from a muscle-like representation (where movements engaging non-overlapping muscles would evoke non-overlapping neural activity).

A modular neural code might be useful for transferring motor skills to different limbs. A motor skill could be learned in movement-coding neural dimensions and then transferred to a different limb by changing the activity in limb-coding dimensions (schematized in Fig. 7). Behaviorally, motor skills such as sequences, rhythms and trajectories have been demonstrated to transfer from one arm to the other (Latash 1999; Criscimagna-Hemminger et al. 2003; Shea, Kovacs, and Panzer 2011), from one leg to the other (Morris et al. 2009), and from the arms to the legs (Kelso and Zanone 2002; Savin and Morton 2008; Christou and Rodriguez 2008). We hypothesize that neurons in the precentral gyrus might help to enable skill transfer by transforming a modular representation of movement inherited from upstream areas into a more muscle-like representation required for motor control. Such an “in-between” role could explain why neural activity in precentral gyrus was not fully modular in the sense that correlations values were always less than one (they typically ranged from 0.5 to 0.7), meaning that matching movements of different limbs could still be distinguished from each other neurally even in movement-coding dimensions alone.

**Figure 7.**
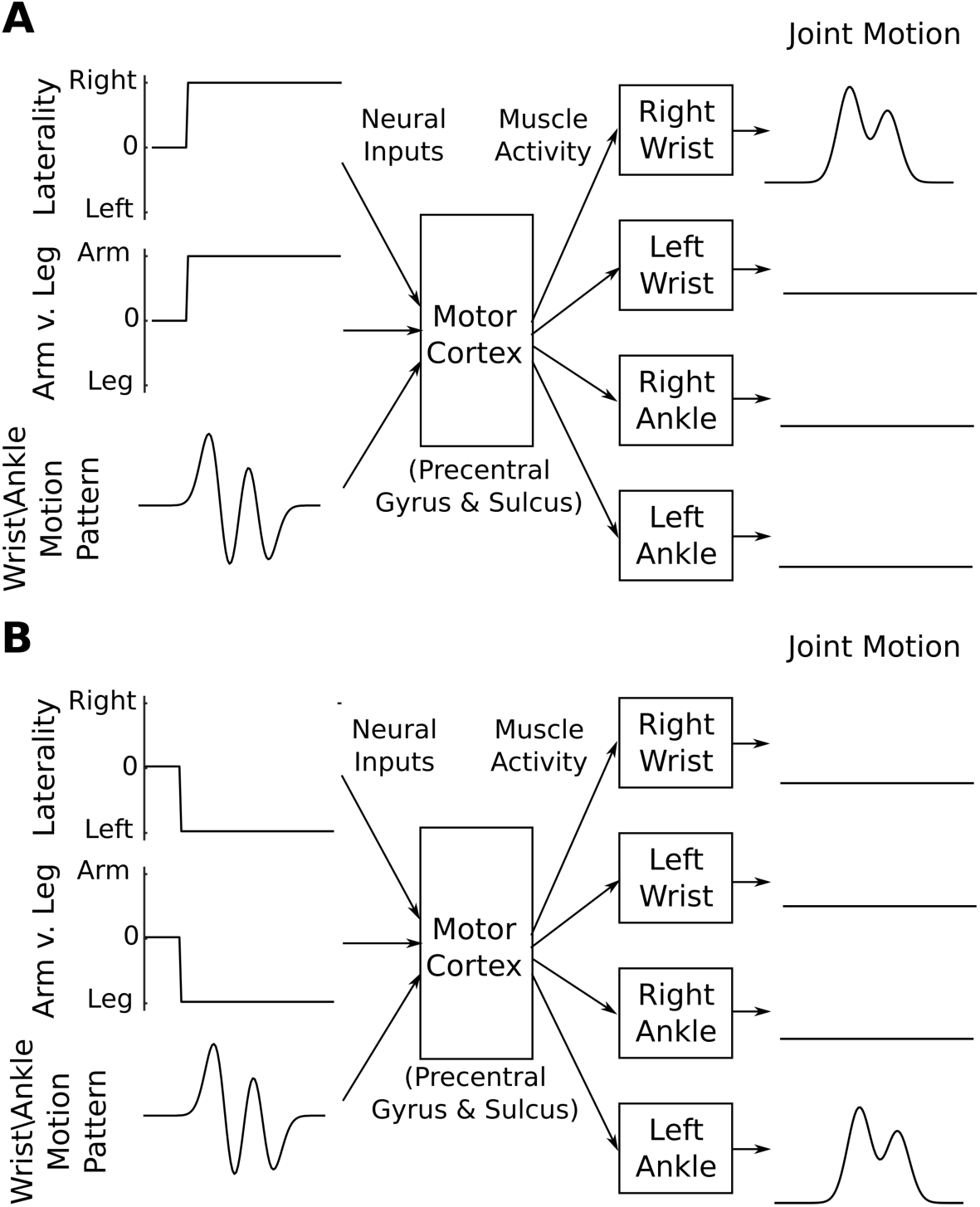
A modular neural code may facilitate the transfer of motor skills across limbs. (A) We speculate that precentral gyrus might receive input from upstream areas in the form of a “modular” representation of movement where the limb is specified somewhat independently of the motion details. In this example, activity in the “laterality” and “arm vs. leg” input streams specify which of the four limbs should move, and activity in the “wrist/ankle motion pattern” input stream specifies the motion that should be performed (at a wrist or ankle joint). Motor areas in the precentral gyrus and central sulcus would then convert this representation into limb-specific muscle patterns to achieve a desired joint motion. (B) The same motion pattern could be transferred to any other wrist or ankle by changing activity only in the “laterality” and “arm vs. leg” dimensions, achieving immediate transfer without requiring relearning.

Our findings are consistent with previous fMRI and ECoG studies that found a correlated representation of ipsilateral and contralateral finger/arm movements on precentral gyrus(Diedrichsen, Wiestler, and Krakauer 2013; Jin et al. 2016; Bundy et al. 2018) and an intrinsic, effector-independent code for finger movements(Wiestler, Waters-Metenier, and Diedrichsen 2014). A recent microelectrode study of human parietal cortex also found that the representation of ipsilateral and contralateral hand and shoulder movements was correlated (Zhang et al. 2017). Our study extends these results to a wider range of movements, to homologous movements of the arms and legs, and to motor cortex. We also show that neural activity in movement dimensions can be prepared independently of neural activity in limb-coding dimensions, lending more support to the idea of independent specification of the movement and the limb to be moved.

We found that when two effectors were moved at the same time, the neural representation for the secondary effector was suppressed and nonlinearly changed, becoming less aligned with the primary effector. These neural changes might facilitate simultaneous control of multiple effectors by helping to keep neural commands related to the secondary effector separate from those of the primary effector. Our results are consistent with prior behavioral work suggesting that bimanual movements are coded separately from unimanual movements (Nozaki, Kurtzer, and Scott 2006; Yokoi, Bai, and Diedrichsen 2016) and prior fMRI studies in people, and electrode recordings in macaques, showing similar nonlinear changes in neural activity (Rokni et al. 2003; Diedrichsen, Wiestler, and Krakauer 2013; Ifft et al. 2013). This study extends these findings to a broader set of effector pairings, showing that there is a rank ordering of effectors that determines which effector’s representation remains intact during simultaneous movement. This finding suggests another aspect to somatotopy on the precentral gyrus: each cortical area may have a preference in terms of which effector’s activity is most conserved during simultaneous movement.

Since the macaque is an important model for the human motor cortical system, it is useful to consider how these results relate to previous findings on macaque motor cortex. It is generally believed that Brodmann’s area 4 (which lies within the central sulcus) is homologous to primary motor cortex in macaques, while Brodmann’s area 6 (which lies on the precentral gyrus) is homologous to macaque premotor areas. Our microelectrode arrays were implanted in dorsal precentral gyrus (hand knob), which has been proposed to be homologous to macaque area F2 (PMd) (Zilles et al. 1995; Rizzolatti, Luppino, and Matelli 1998; Geyer et al. 2000; Genon et al. 2017, 2018). From this perspective, many of our results match previous results from macaque studies, including a large correlation in PMd activity between contralateral and ipsilateral reaching movements (but not in M1) (Cisek, Crammond, and Kalaska 2003; Ames and Churchland 2019; Heming et al. 2019), and mixed arm and leg representations in PMd (but not in M1) (Kurata 1989). In Supplemental Figure 9, we directly compare our data to two recent studies on macaque M1(Ames and Churchland 2019; Heming et al. 2019) and find that there are substantially higher correlations between ipsilateral and contralateral movements in our data. It is possible that, had we been recording deep in the central sulcus, we would have seen sharper somatotopy and weaker correlations.

It is important to keep in mind that both participants had chronic tetraplegia (T7 had ALS and T5 had high level spinal cord injury), which raises the question of how these findings might generalize to people without tetraplegia. One would expect cortical remapping to occur in the case of spinal cord injury; however, current models of remapping predict that the immediately adjacent intact body parts (in this case face and head) should become more dominant (Qi, Stepniewska, and Kaas 2000). Instead, we found in participant T5 that arm and leg movements were more strongly represented than face and head movements, suggesting that the extent of remapping may be limited in precentral gyrus; consistent with this, a recent fMRI study also found limited motor cortical remapping in hand amputees (Wesselink et al. 2019). The fact that we observed roughly the same proportions of modulation for face, head, arm and leg movements in participant T7, who had ALS, gives further confidence that the broad tuning we observed was not caused by remapping. While there is always the possibility that any of our findings may have been caused by some unknown process of cortical change due to spinal cord injury or ALS, we know of no specific reason to believe this to be the case.

Finally, these results have important implications for intracortical brain-computer interfaces (iBCIs), since it opens up the opportunity to decode movements across the entire body from just a small area of precentral gyrus. We showed, for the first time, that a discrete target selection iBCI could successfully decode targets across all four limbs accurately enough to improve performance relative to a single-effector approach. Other use cases could be to map multiple movements to different kinds of clicks for general purpose computer use (e.g. curl left toes for left click, curl right toes for right click, point foot up to scroll up, etc.), or even to restore continuous control of leg and arm movements across both sides of the body. Our findings on the neural representation of single effector and dual-effector movements may help to inform the design of advanced decoders that enable control over multiple body parts.

## Author Contributions

F.R.W. conceived the study and wrote the manuscript. F.R.W., D.R.D., D.T.A, and P.R. collected the data. D.R.D. took the lead on developing the closed-loop discrete decoding system and related experiments (Fig. 6). F.R.W. took the lead on designing, analyzing, and interpreting all other experiments. L.R.H. is the sponsor-investigator of the multi-site clinical trial. J.M.H. planned and performed T5’s array placement surgery and was responsible for his ongoing clinical care. J.M.H. and K.V.S. supervised and guided the study. All authors reviewed and edited the manuscript.

## Acknowledgements

We thank participants T5 and T7 and their caregivers for their dedicated contributions to this research, N. Lam for administrative support, and J. Simeral and B. Sorice for data collection with T7.

This work was supported by the A. P. Giannini Foundation, US NINDS Transformative Research Award R01NS076460, NIH Director’s Pioneer Award 8DP1HD075623-04, Director’s Transformative Research Award (TR01) from the NIMH #5R01MH09964703, NIDCD R01DC014034, NINDS R01NS066311, NIDCD R01DC009899, NICHD-NCMRR N01HD53403 and N01HD10018; DARPA REPAIR N66001-10-C-2010; Department of Veterans Affairs Rehabilitation Research and Development Service B6453R; MGH-Deane Institute for Integrated Research on Atrial Fibrillation and Stroke; Simons Foundation.

## Methods

### Study Permissions and Participants

This study includes data from two participants (identified as T5 and T7), who gave informed consent and were enrolled in the BrainGate2 Neural Interface System clinical trial (ClinicalTrials.gov Identifier: NCT00912041, registered June 3, 2009). This pilot clinical trial was approved under an Investigational Device Exemption (IDE) by the US Food and Drug Administration (Investigational Device Exemption #G090003). Permission was also granted by the Institutional Review Boards of University Hospitals (protocol #04-12-17), Stanford University (protocol #20804), Partners Healthcare/Massachusetts General Hospital (2011P001036), Providence VA Medical Center (2011-009), and Brown University (0809992560). All research was performed in accordance with relevant guidelines/regulations.

Participant T7 was a right-handed man, 53 years old at the time of data collection, who was diagnosed with Amyotrophic Lateral Sclerosis (ALS) and had resultant motor impairment (functional rating scale ALSFRS-R of 17). In July 2013, T7 was implanted with two 96 channel intracortical microelectrode arrays (Blackrock Microsystems, Salt Lake City, UT) in the hand knob area of dominant precentral gyrus (1.5-mm length). Data are reported from post-implant day 24. At the time of data collection, T7 retained movement of the arm, leg, face and head. T7 was able to extend and flex the knee and ankle normally, and make all requested head and face movements normally, but had more limited range of motion in the arm for some movements. T7 is no longer enrolled in the trial and the data from T7 were collected before the present study was conceived.

Participant T5 is a right-handed man, 65 years old at the time of data collection, with a C4 ASIA C spinal cord injury that occurred approximately 9 years prior to study enrollment. In August 2016, participant T5 was implanted with two 96 channel intracortical microelectrode arrays (Blackrock Microsystems, Salt Lake City, UT) in the hand knob area of dominant precentral gyrus (1.5-mm length). Data are reported from post-implant days 579-961 (Supplemental Table 1). T5 retained full movement of the head and face and the ability to shrug his shoulders. Below the injury, T5 retained some very limited voluntary motion of the arms and legs that was largely restricted to the left elbow. Some micromotions of the right and left hands were also visible, along with extremely small but reliable twitching of the feet and toes.

The hand knob area in both participants was identified by pre-operative magnetic resonance imaging (MRI). Supplemental Figure 2 shows array placement locations registered to MRI-derived brain anatomy.

### Session Structure and Cued Movement Tasks

Neural data was recorded in 3-5 hour “sessions” on scheduled days. During the sessions, participants sat upright in a wheelchair that supported their backs and legs. A computer monitor placed in front of the participants displayed text and/or colored shapes to indicate which movement to make and when. Data was collected in a series of 2-10 minute “blocks” consisting of an uninterrupted series of cued movements; in between these blocks, participants were encouraged to rest as needed.

The data from participant T7 were collected in a single session consisting of a series of 2-minute blocks containing 20 repetitions of one set of paired movements. These paired movements were designed to engage the same joint(s) but in different directions; for example: hand open/close, wrist flexion/extension, head left/right, etc. Every three seconds, the text on the screen alternated to instruct the other paired movement (e.g. as in Fig 1A) and T7 responded as soon as he was able.

The data from T5 were collected across 17 sessions (Supplemental Table 1). All sessions followed a simple instructed delay paradigm (like that shown in Fig 1A) except for the closed-loop discrete decoding sessions (described in their own section below). For text-cued tasks (Figures 1,2,3 & 5), during the instructed delay period a red square and text appeared in the center of the screen indicating to T5 that he should prepare to make the specified movement. The instructed delay period lasted a random amount of time that was drawn from an exponential distribution; values that fell outside of a minimum/maximum range were re-drawn. Maximum and minimum delay times varied from session to session but were within 1.4 to 3 seconds. After the delay time, the square turned green and the text indicating the movement changed to “Go”, at which point T5 made the movement immediately. T5 was told to continue attempting to make the movement (for arm & leg movements) or to hold the posture of the completed movement (for face & head movements) until the text changed to “Return”, at which point T5 relaxed and returned to a neutral posture. The movement and return periods lasted 1.5 seconds each. The spatially cued movement task (Fig 4) followed the same structure, but used colored targets instead of text to specify which movement was supposed to be made (as shown in Fig 4A).

### Neural Data Processing

Neural signals were recorded from the microelectrode arrays using the NeuroPort(tm) system (Blackrock Microsystems) [(Hochberg 2006) describes the basic setup]. Neural signals were analog filtered from 0.3 Hz to 7.5 kHz and digitized at 30 kHz (250 nV resolution). Next, a common average reference filter was applied that subtracted the average signal across the array from every electrode in order to reduce common mode noise. Finally, a digital bandpass filter from 250 to 3000 Hz was applied to each electrode before spike detection.

For threshold crossing detection, we used a -4.5 x RMS threshold applied to each electrode, where RMS is the electrode-specific root mean square (standard deviation) of the voltage time series recorded on that electrode. For the analysis of waveform-sorted single neurons (Supplemental Figure 3), neurons were sorted manually using Plexon Offline Spike Sorter v3. Only waveform clusters that appeared in an unambiguous, highly separable cluster in principal component space were included.

Threshold crossing times (and single-unit spike times) were binned into 10 ms bins (for T5) or 20 ms bins (for T7) and z-scored to ensure that electrodes with high firing rates did not overly influence the population-level results. Different bin widths were used for T5 and T7 because the task data was collected with different computer systems at different sampling rates. Z-scoring was accomplished by first subtracting, in a block-wise manner, the mean count over all bins within each block. Subtracting the mean within each block helps to counteract slow drifts in the firing rate of electrodes and was especially important for the T7 data, since it was collected in a block-wise fashion. For this dataset, mean subtraction ensures that spurious drifts in firing rate did not artificially inflate the classification performance shown in Fig 1D (by helping to distinguish movements from different blocks). After mean-subtraction, the binned counts were divided by the sample standard deviation computed using all bins across all blocks.

Finally, electrodes with firing rates < 1 HZ over all time steps were excluded from further analysis in order to denoise population-level results. For some analyses (e.g. dPCA in Fig 2B), the binned spike count time series were convolved with a Gaussian kernel (sd = 30 ms) to reduce high frequency noise.

### Gaussian Naïve Bayes Classification

For both offline (e.g. Fig. 1D) and online classification (Fig. 6 F-G), we used a Gaussian naïve Bayes classifier to classify firing rate vectors. Firing rate vectors were computed by counting the number of threshold crossings that occurred within a fixed window of time for each electrode and then dividing by the window width.

To classify the firing rate vectors, we computed the log likelihood of observing that vector for each possible class and then chose the class with the largest log likelihood. We assumed that each element of the firing rate vector was normally distributed and was statistically independent from each other element. We also assumed that the variance of each Gaussian distribution was the same across all classes. Under these assumptions, the following quantity is proportional to the log likelihood of observing firing rate vector ***x*** under the class *C*_*k*_:

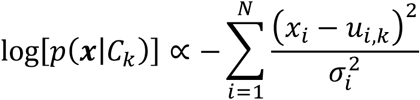

Here, N is the number of electrodes, *x*_2_ is the *i*th element of the firing rate vector **x** (corresponding to electrode *i*’s firing rate), *u*_*i,k*_ is the mean for electrode *i* under class *k*, and σ_i_ is the variance for electrode *i* (under all classes). To classify, we computed this quantity for each class and chose the class with the largest value. The class-specific means were estimated using the sample mean across all available training data; likewise, the variances were estimated using the pooled variances across all classes.

### Closed-Loop Discrete Decoding

All closed-loop discrete decoding experiments used a “time-locked” trial structure where each trial lasted for 1 second and no pauses occurred between trials. At the end of each trial, the closed-loop discrete decoding system classified the participant’s attempted movement. If the movement was correctly classified, a “success” sound played; otherwise, a “failure” sound played. After each trial ended, a new target appeared and the next trial began immediately.

We used a Gaussian naïve Bayes classifier (defined above) to classify mean firing rate vectors computed using a fixed window of time for each trial. This window was optimized for each task and session as part of the process of calibrating the classifier; the window typically fell between 300 to 1000 ms. To optimize the window, we performed an offline grid search across all possible start and end times for the window in 50 ms increments from 0 ms to 1000 ms. Ten-fold cross-validation was used to estimate the achieved bit rate that would result from using each window. The window with the largest achieved bit rate was then chosen.

For each online discrete decoding session, one or two target configurations were tested. For each target configuration, one 5 minute open-loop block of data was first collected with which to calibrate an initial classifier. Then, we proceeded with a series of 5 minute closed-loop blocks to allow participant T5 to practice with that target configuration. After each closed-loop block, we re-calibrated the classifier using the preceding two blocks of data. We continued until we determined that T5’s performance had plateaued and he was comfortable with the task. After this decision was made, we collected a series of five, 3-minute “performance” blocks with the classifier held fixed; the average accuracy and achieved bit rate obtained during these blocks were reported in Fig. 6.

Achieved bit rate is a conservative, clinically-motivated metric of information throughput (Nuyujukian 2015) and was computed with the following formula:

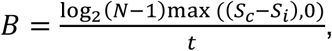

where *N* is the number of targets, *S*_*c*_ is the number of correct selections, *S*_*i*_ is the number of incorrect selections, and *t* is the total time elapsed. Achieved bit rate assumes that every incorrect selection must be followed by the correct selection of a delete key to undo the error, thus heavily penalizing low accuracies that would be hard for participants to work with in practice.

For the online results in Fig 6 F-G, we used an offline simulation to find the optimal number of targets for each layout in order to maximize the achieved bit rate. To do so, we used an exploratory offline dataset with all four effectors (right & left hand, right & left foot) and with eight targets per effector. We then fit a simple linear encoding model of target direction to each effector, using only the data corresponding to that effector’s movements:

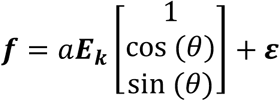

Here, *f* is the N × 1 firing rate vector for a single trial, ***E***_***k***_ is an N ×3 matrix of mean firing rates (first column) and directional tuning coefficients (second and third columns) for effector *k*, θ is the angle of the target, ***ε*** is an N ×1 noise vector, and *a* is a scalar gain factor. Each element of ***ε*** is normally distributed with covariance matrix **Σ**_**k**_. ***E***_***k***_ was fit to the data using least squares regression for each effector independently, and **Σ**_**k**_ was estimated using the sample error covariance of the linear model. To counteract overfitting in ***E***_***k***_ (which could overestimate the amount of tuning present), we searched for the gain factor *a* which led to the best match in predicted accuracy for the number of targets present in the exploratory dataset (eight). These *a* values were < 1 and downweighed the tuning.

Once the model was fit, we varied the number of possible targets from 2 to 10. For each target layout, we simulated a new set of trials using the model and predicted decoder performance by applying a Gaussian naïve Bayes classifier to the simulated data using ten-fold cross-validation. Using this method, we predicted that the optimal number of targets was equal to 6 for a single effector (right hand only), 6 each for two effectors (right & left hand), and 4 each for four effectors (right & left hand, right & left foot) when the trial time was constrained to be one second. Encouragingly, the optimal number of targets predicted by this method for a single effector (6) matched online data (Fig 6b).

### Median-Unbiased Measurements of Firing Rate Distance

Measuring the distance between two distributions of firing rate vectors is useful for quantifying the amount of neural modulation caused by a particular movement. However, measuring this in an unbiased way in the presence of noise is a nontrivial problem. Here, we motivate the problem and describe the methods we used to obtain median-unbiased estimates of distance by using cross-validation (see our code repository at https://github.com/fwillett/cvVectorStats for an implementation).

To understand the problem, consider the measurements in Fig 1C. The bar heights reflect the distance between two multivariate distributions: a distribution of firing rate vectors corresponding to the “do nothing” condition (distribution 1), and a distribution of firing rate vectors corresponding to the movement in question (distribution 2). Let us denote the firing rate vector observed on trial *i* for distribution 1 as 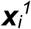 and for distribution 2 as 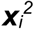. The sample means are

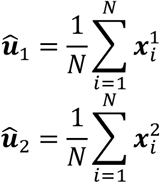

The sample means differ from the true means *u*_1_ and *u*_2_ by some sampling error. As a result, the difference in sample means also contains error: 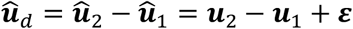 for some error vector ***ε***.

One simple method for estimating the distance between distributions 1 and 2 would be to take the Euclidean norm of the difference in sample means: 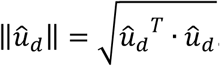. However, the sampling error inherent in *û*_*d*_ causes this metric to be biased upwards. Expanding terms, we have: 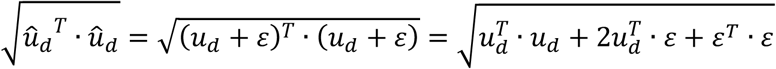, which is larger than 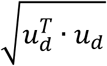 on average since *ε*^*T*^ · *ε* > 0. One intuitive way to see this is to consider the case where distribution 1 and distribution 2 are identical. The sample means for each distribution will always differ from each other (due to sampling error), and hence there will always be some positive distance between them even though the true difference in means is zero.

Fortunately, there is a straightforward way to achieve a median-unbiased estimate of distance. First, by following the approach laid out in (Allefeld and Haynes 2014), we can estimate the squared distance by averaging together a series of unbiased estimates, each made by using leave-one-out trials. The following is an unbiased estimate of the squared distance: 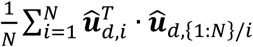, where 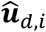 is the sample estimate of the difference in means using only the pair of samples from trial *i* and 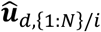 is the sample estimate of the difference in means using all trials except *i*. Since these two estimates contain no overlapping trials, they are statistically independent from each other. Statistical independence implies that the expectation of their product is equal to the product of their expectation. Taking the expectation, we can show that it is an unbiased measure:

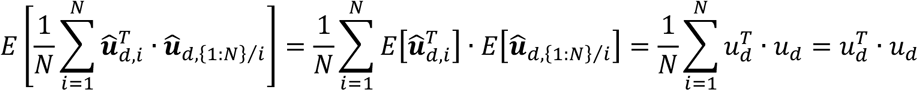

Note that this cross-validation approach will work for computing the squared norm of any vector quantity in an unbiased way. For example, 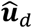 could be a vector of coefficients from a linear tuning model or, more simply, the mean of a single distribution.

Unfortunately, taking the square root of this quantity does not immediately yield an unbiased measure of distance, since this quantity can now be negative. It is possible to construct a simple, median-unbiased measure by taking the square root of the absolute value while preserving the sign: 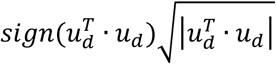 Since this function is a monotonic function of an unbiased estimate, it is median-unbiased; however, since the square root is nonlinear and concave, this estimate is not mean-unbiased and is conservative (in cases of high sampling error, it will be smaller than the true value). Nevertheless, it is still far more robust to sampling error than the naïve approach of taking the Euclidean norm of the sample estimate. In simulation, we found that except in cases of very high sampling error, its expectation was equal to the true value unlike the naïve approach (Supplemental Figure 10).

Note that above method assumes that there are an equal number of observations from each distribution (N observations from each), but the same technique can be applied for unequal numbers of observations by splitting the data into folds of varying sizes (https://github.com/fwillett/cvVectorStats).

### Reducing Bias when Estimating Vector Angles and Correlations

In the above section, we described our approach for computing median-unbiased estimates of the distance between two firing rate distributions. We extended this approach to reduce the bias of correlation and angle measurements, which we describe here (see our code repository at https://github.com/fwillett/cvVectorStats for an implementation).

When computing the angle between mean firing rate vectors (in Fig 5F), the sample estimate is:

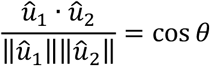

Here, *û*_1_ and *û*_2_ are the sample estimates of the mean firing rate vectors for distribution 1 and 2. The problem here is that ‖*û*_1_‖ and ‖*û*_2_‖ are biased upwards because they contain sampling error. This causes cos *θ* to appear smaller than it truly is, making the vectors appear more dissimilar. The larger the uncertainty (smaller sample sizes or smaller vectors), the greater this effect will be.

By replacing ‖*û*_1_‖ and ‖*û*_2_‖ with median-unbiased estimators of vector norm, this effect can be eliminated. For example, as shown in the above section, ‖*û*_1_‖^2^ can be estimated in an unbiased way by computing 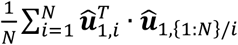, where 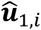 is the sample estimate of ***u***_1_ using only trial *i* and 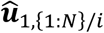 is the sample estimate of ***u***_1_ using all trials except *i*. ‖*û*_1_‖ can then be estimated by applying the sign-preserving square root transform.

The same technique of substituting median-unbiased estimators into the denominator can be applied to reduce bias when estimating correlations between vector means (as in Fig 3) or vectors of tuning coefficients (as in Fig 4). To see this, note that the equation 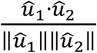 estimates the correlation between two vectors *u*_1_ and *u*_2_ as long as *û*_1_ and *û*_2_ are taken to be the sample estimates of the *centered* vectors. In Supplemental Figure 10, we show that this approach substantially reduces bias and practically eliminates it compared to a naïve approach.

In Fig 1C, we used the median-unbiased estimator of distance to compute the height of each bar. To compute the confidence intervals, we used a jackknife procedure as described in (Severiano et al. 2011). We found that the jackknife confidence intervals produced more accurate confidence intervals than bootstrapping (which we confirmed via simulation), since the cross-validated distance metric is biased upwards when resampling identical data points (because identical data points are more aligned with each other). This effect is similar to that noted in (Severiano et al. 2011), where the jackknife was also found to outperform the bootstrap.

To compute the small circles in Fig 1C that represent single trial firing rates, we used cross-validation to project each trial’s firing rate vector onto a line connecting the “do nothing” distribution to the movement distribution in question. For each trial, we first estimated the line connecting the two distributions using all other trials by computing *ν* = (*û*_2_ − *û*_1_)/*d*, where *û*_1_ is the sample mean for distribution 1, *û*_2_ is the sample mean for distribution 2, and *d* is the scalar distance between the distributions found using the median-unbiased estimator. When the projected the held-out trial’s firing rate vector onto *ν* by using the dot product.

For Fig 1D (offline classification), we used the firing rates in a window between 200-600 ms (T5) or 200-1600 ms (T7) after the go cue as input into the classifier. For T7, we found that since there was temporal variation in activity throughout the movement, classification performance could be improved by including multiple time windows of firing rates for each electrode in the feature vector. We split the 200 to 1600 ms time window into two smaller windows (200 to 800 and 800 to 1600) and included the activity in each of these windows for each electrode as input to the classifier (effectively doubling the number of inputs). This wasn’t necessary for the T5 dataset as performance was already saturated with a single window.

Fig 2A was created in the same way as Fig 1C. For Fig 2B-D, we used the most recent version of demixed principal components analysis (Kobak et al. 2016) [https://github.com/machenslab/dPCA]. Demixed principal components analysis (dPCA) is an optimization technique that decomposes neural data into a sum of neural dimensions that express variance related only to certain variables in the experiment. This decomposition gives an interpretable and comprehensive overview of the structure in the neural data as it relates to the experimentally manipulated variables. Here, we briefly summarize dPCA as it was applied to our data [refer to (Kobak et al. 2016) for a complete description].

Demixed PCA begins with splitting the data into marginalizations in an ANOVA-like manner. Considering first just a single electrode, we can put this electrode’s binned firing rate data into a four-dimensional data tensor *x*_tmlk_, where *t* is the time step, *m* is the movement condition, *l* is the laterality condition, and *k* is the trial. By averaging across trials, we can obtain the trial-averaged, three-dimensional data tensor 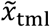.

We can now define the following marginalizations of the trial-averaged data tensor. In the following equations, when a subscripted index is replaced with a dot, it means to average over that dimension:

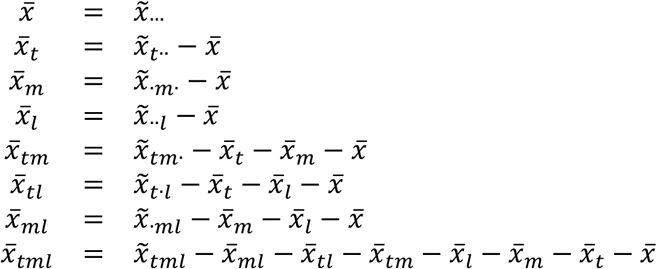

The left-hand side of the above equations are the marginalizations and the right-hand side shows how to compute them. Note that the trial-averaged data tensor can be written as a sum of the marginalizations, since they define a complete decomposition of the data:

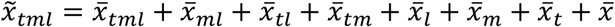

Next, each marginalization can be “unrolled” into a row-vector of length TML by tiling the marginalization appropriately (where T is the number of time steps, M is the number of movement conditions, and L=2 is the number of body sides). Once unrolled, this marginalization can be “stacked” vertically across all electrodes to yield a matrix of size N x TML that describes the entire neural population’s marginalization.

After computing the marginalizations, demixed PCA then attempts to find neural dimensions that “readout” certain sets of the marginalizations on a single-trial basis. For example, we can group together the 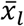 and 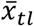 marginalizations by adding together their matrices *X*_*L*_ and *X*_*TL*_, yielding an N x TML matrix of laterality-related signals in the neural population. Demixed PCA then tries to find neural dimensions that will map single trial recordings to the marginalization matrix. To do so, first the marginalization matrix must be expanded to tile across all single trials, yielding a matrix of size N x TMLK (where K is the number of trials). Demixed PCA then uses the following loss function:

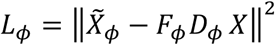

Here, 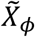 is the trial-averaged marginalization set of interest (in this example, *X*_*L*_ + *X*_*TL*_) tiled across all trials, X is a matrix containing the single-trial data (also of size N x TMLK), and D and F are decoder and encoder matrices (of size C x N and N x C) that readout the single trial activity into C signals and then re-encode them back into the N-dimensional neural activity.

For the analysis in Fig 2B, we grouped the marginalizations as follows for ease of interpretation: movement 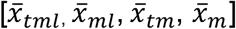, laterality 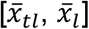, and time.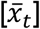 Fig 2B shows the top readout dimensions (rows of *D*_*ϕ*_ *X* in order of variance explained) for each marginalization group. Each line depicts the average across trials and the shaded regions depict 95% confidence intervals. Confidence intervals were found using a cross-validated version of dPCA. Specifically, for each trial, we performed dPCA on all remaining trials to find the readout dimensions *D*_*ϕ*_ and then applied them to the held-out trial. After doing this for each held-out trial, we then estimated the means and 95% CIs using the held-out readouts. Cross-validation ensures that any result found is not due to overfitting the decoder matrices to noise in the data. One potential issue with this cross-validation approach is that the sign of the readout dimensions can sometimes change for each held-out trial; to counteract this, we corrected for any sign flips by finding the sign that best matched the readout dimensions found for the first held-out trial (match quality was assessed with the dot product). This problem is discussed in general in (Milan and Whittaker 1995).

In Fig3A, we used a cross-validation approach to verify that the laterality and arm vs. leg dimensions found by dPCA generalized to new movements. To do so, we took two approaches: a held-out “single movement” approach and a held-out “movement set” approach. In the single movement approach (diagonal panels in Fig 3A), we used leave-one-out cross-validation for each single movement condition. That is, for each movement, we applied dPCA to all other movements in order to find the top component for reading out the laterality or arm vs. leg marginalization. We then applied this component to the held-out condition. In the movement set approach (off-diagonal panels in Fig 3A), we held out an entire set of movements corresponding to a different limb (or laterality). The results show that the laterality and arm vs. leg components generalize both across movement conditions and across different sides of the body and effectors.

For Fig 3B-C we used the bias-corrected correlation metric (described above in “Reducing Bias when Estimating Vector Angles and Correlations”) to compute the correlation between pairs of mean firing rate vectors associated with each movement condition. Fig 3B uses data collected in session 12.05.2018 (the same as in Fig2), while Fig3C used a different dataset designed specifically to contain many pairs of homologous arm and leg movements (11.19.2018). Before computing the correlations, we subtracted the average firing rate within each laterality (Fig 3B) or within each effector (Fig 3C). Otherwise, correlations would be predominately negative across all movements due to the presence of large laterality and arm vs. leg dimensions (shown in Fig 3A).

Fig 3D contains data from two separate sessions (12.10.2018 and 03.27.2019) designed to test whether partially specified movements could elicit preparatory activity in dimensions corresponding only to the partially specified information. In the partial laterality cueing experiment (top panels), there were four movements tested: contralateral wrist extension, ipsilateral wrist extension, contralateral hand grasp, and ipsilateral hand grasp. During the instructed delay period, the upcoming movement could either be fully cued (with both the laterality & movement specified by text on the screen), laterality-cued (with only the laterality specified), or movement-cued (with only the movement specified), with equal probability of each. In the partially cued conditions, when the go cue was given, the text on the screen changed to specify the missing piece of information so that the movement could be attempted.

For example, for a movement-cued trial for contralateral wrist extension, during the delay period a red square appeared in the center of the screen and the text above it was: “Wrist”. Then, during the go period, the square turned green and the text changed to: “Right”. Displaying “Right” instead of “Right Wrist” during the go period ensured that T5 must remember the partially specified piece of information (and cannot simply skip movement preparation, waiting until the go cue to read the text).

The partial arm vs. leg cueing experiment (bottom panels of Fig 3D) had an analogous design with the following four movements: wrist up (extension), wrist down (flexion), ankle up (dorsiflexion), and ankle down (plantarflexion). During the instructed delay period, the upcoming movement could either be fully cued (with both the effector & movement direction specified by text on the screen), effector-cued (with only the “wrist” or “ankle” specified) or movement-cued (with only the movement direction specified).

To analyze the data from the partial cue experiment, the data was first smoothed (convolved with a Gaussian kernel, width = 60 ms std) and marginalized according to either movement type or laterality (for the first experiment), or movement direction or effector (for the second experiment). We then applied demixed PCA to a time window of the data beginning 1 second before the go cue and ending at the go cue (see the section above on Fig 2 for an explanation of dPCA and marginalization). The demixed PCA components were found using only the fully cued trials.

To make Fig 3D (and Supplemental Figure 6), we used the dPCA component for each marginalization that explained the most variance. The means and confidence intervals plotted in Fig 3D were found by first averaging the activity in each dPCA dimension across a 300 ms time window for each trial. This yields a single value for each trial; these values were then averaged for each condition and confidence intervals were fit by assuming a normal distribution. We used unsmoothed neural data for this portion of the analysis so that only data that was strictly contained within the 300 ms window was used.

To make Fig 4B, we used PCA to find, for each effector separately, the two highest-variance neural dimensions that best explained across-target variance. To do so, we first computed a firing rate vector for each trial containing each electrode’s firing rate within a window from 200 to 1000 ms after the go cue. Next, we averaged these firing rate vectors within each target to obtain eight target-averaged vectors. We then applied PCA to these target-averaged vectors to obtain the top two neural dimensions that best explained across-target variance. Finally, we then projected each trial’s firing rate vector onto these components to obtain the single-trial dots shown in Fig 4B. These were then rotated using a Procrustes analysis to align with the target geometry (i.e. a single rotation matrix was found that best mapped the neural activity to the target locations; this matrix had no ability to shear or skew the firing rates).

We used cross-validation to ensure that the result in Fig 4B was not due to overfitting noise in the data. When applying PCA, we used a leave-one-out cross-validation approach. For each trial, the top two neural dimensions were found using all other trials; the held-out trial was then projected onto those neural dimensions. One potential issue with a cross-validation approach for PCA is that the sign of the PCs can sometimes change for each held-out trial; to counteract this, we corrected for any sign flips by finding the sign that best matched the PCA dimensions found for the first held-out trial (match quality was assessed with the dot product). This problem is discussed in general in (Milan and Whittaker 1995).

When computing the correlation between vectors of linear tuning coefficients, we used the bias-reduced metrics described above in “Reducing Bias when Estimating Vector Angles and Correlations”. Tuning coefficients were found using the following model:

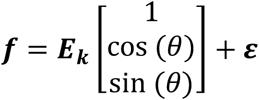

Here, ***f*** is the N × 1 firing rate vector for a single trial, ***E***_***k***_ is an N x 3 matrix of mean firing rates (first column) and directional tuning coefficients (second and third columns) for effector *k*, θ is the angle of the target, and ***ε*** is an N × 1 noise vector. ***E***_***k***_ was fit using ordinary least squares regression using all trials from effector *k*. Correlations were computed by calculating Pearson’s r using matching columns of ***E***_***k***_ for pairs different effectors.

To find the neural dimensions used to visualize the activity in Fig 5B-D, we first computed a firing rate vector for each trial containing each electrode’s firing rate within a window from 200 to 1000 ms after the go cue. For Fig 5B, in order to find neural dimensions for visualization, we first averaged these firing rate vectors within each single-movement condition to obtain four average vectors: **f**_**1A**_, **f**_**1B**_, **f**_**2A**_, **f**_**2B**_ (for effectors 1 & 2 and movement directions A & B). We then subtracted the average firing rate vector within each effector to remove the differences in firing rates caused by effector alone. That is, we computed mean-subtracted vectors:

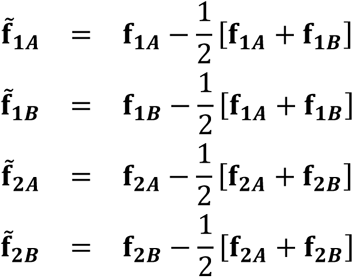

We then applied PCA to the mean-subtracted vectors 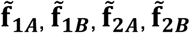 to find the two components that explained the most variance in these vectors. Finally, we used these two components to project the single trial firing rate vectors into a two-dimensional space for visualization. Note that applying PCA to the mean-subtracted vectors ensures that the PCA components capture differences in firing rate that are related to movement direction. Otherwise, PCA would find a large “effector” dimension (like that shown in Fig 3A) and the direction-related activity would not be as visible in the first two dimensions.

For Fig 5D, in order to find the dual movement neural dimensions for visualization, we first averaged the single trial firing rate vectors within each dual-movement condition to obtain four average vectors: **g**_**AA**_, **g**_**AB**_, **g**_**BA**_, **g**_**BB**_ (for movement directions A & B). We then applied PCA directly to these vectors without mean-subtraction. No mean subtraction was necessary in this case because both effectors were active for all movements; thus, there is no possibility of finding dimensions that are related to the effector being moved and not the direction it is moving in.

When applying PCA for Fig 5B-D, we used cross-validation to ensure that the clustering observed was not due to overfitting noise in the data (to cross-validate, we used the same methods as described above for Fig 4).

For Fig 5E and the top panel of Fig 5F (dual/single ratio), we used the median-unbiased estimate of distance (see above sections). For the bottom panel of Fig 5F (change in angle), we used the bias-corrected metric of vector angle (see above sections).

Data for Fig 6B was collected in a single session (11.05.2018) where multiple target layouts were tested. For each target layout, T5 first completed a 4 minute “open-loop” block where he made the cued movements but they were not decoded in real-time. This block was used to calibrated an initial decoder. T5 then completed a 4 minute “practice” closed-loop block using the initial decoder. Next, a new decoder was calibrated by combining data from the open-loop and closed-loop block. Finally, T5 completed a series of three, 4 minute “performance” closed-loop blocks using that decoder; performance form these blocks was reported in Fig 6B.

Data for Fig 6C-E was collected in a separate session (01.07.2019). In this session, T5 completed a series of 15 open-loop blocks with three target layouts (5 blocks each): radial 16 single-effector (right hand joystick), radial 8 dual-effector (left & right hand joystick), and radial 4 quad-effector (left & right hand, left & right feet). Each block lasted 5 minutes. The blocks were interleaved in sets of 3 blocks each (1 for each layout).

Data for Fig 6F was collected in a series of 7 sessions (see above section “Closed-Loop Discrete Decoding” for details about the session structure and decoder). Supplemental Table 1 lists each individual session.

## Supplemental Materials

**Supplemental Table 1.**
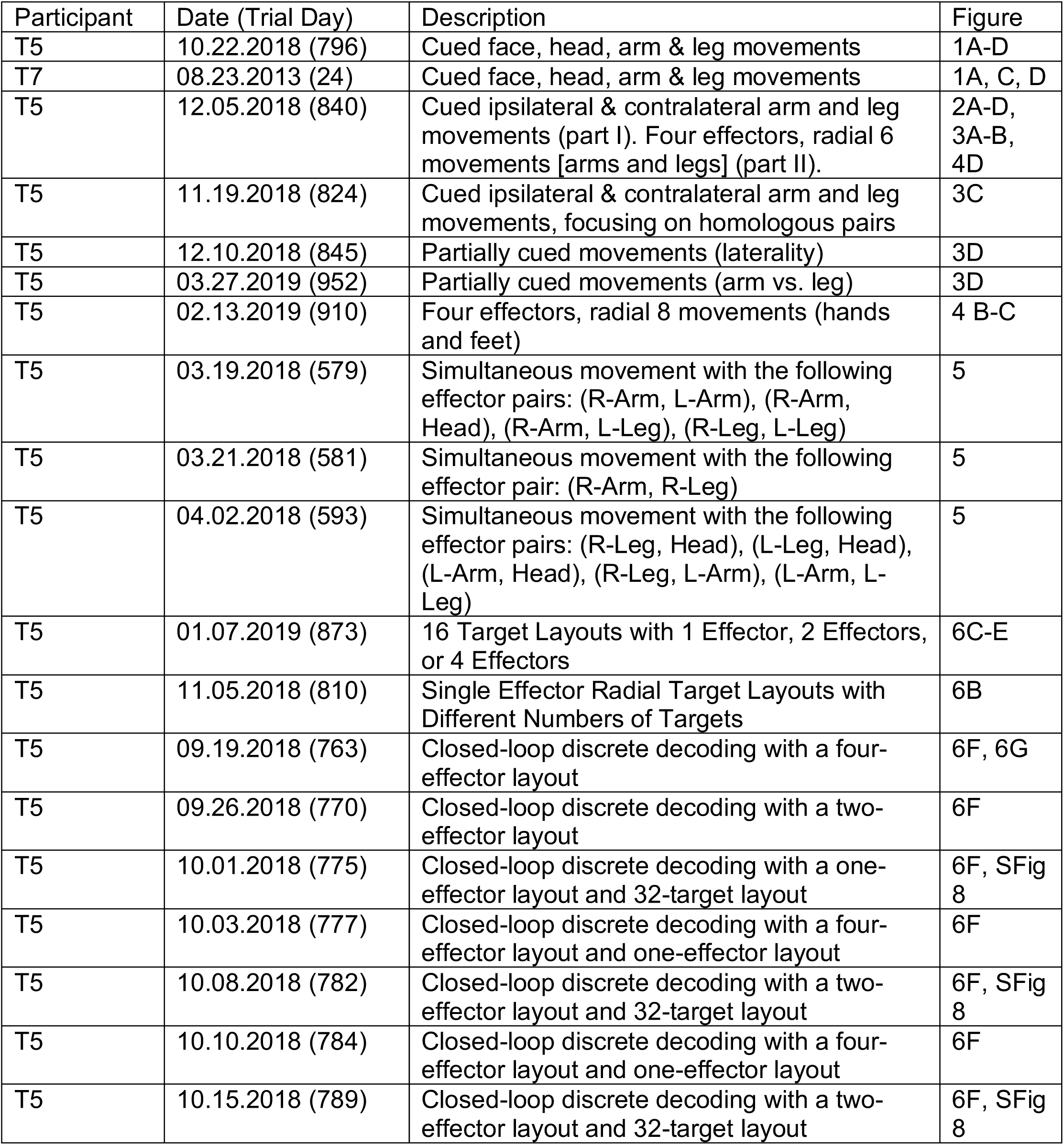
A table of all data collection sessions included in this study.

| Participant | Date (Trial Day) | Description | Figure |
| --- | --- | --- | --- |
| T5 | 10.22.2018 (796) | Cued face, head, arm & leg movements | 1A-D |
| T7 | 08.23.2013 (24) | Cued face, head, arm & leg movements | 1A, C, D |
| T5 | 12.05.2018 (840) | Cued ipsilateral & contralateral arm and leg movements (part I). Four effectors, radial 6 movements [arms and legs] (part II). | 2A-D, 3A-B, 4D |
| T5 | 11.19.2018 (824) | Cued ipsilateral & contralateral arm and leg movements, focusing on homologous pairs | 3C |
| T5 | 12.10.2018 (845) | Partially cued movements (laterality) | 3D |
| T5 | 03.27.2019 (952) | Partially cued movements (arm vs. leg) | 3D |
| T5 | 02.13.2019 (910) | Four effectors, radial 8 movements (hands and feet) | 4 B-C |
| T5 | 03.19.2018 (579) | Simultaneous movement with the following effector pairs: (R-Arm, L-Arm), (R-Arm, Head), (R-Arm, L-Leg), (R-Leg, L-Leg) | 5 |
| T5 | 03.21.2018 (581) | Simultaneous movement with the following effector pair: (R-Arm, R-Leg) | 5 |
| T5 | 04.02.2018 (593) | Simultaneous movement with the following effector pairs: (R-Leg, Head), (L-Leg, Head), (L-Arm, Head), (R-Leg, L-Arm), (L-Arm, L-Leg) | 5 |
| T5 | 01.07.2019 (873) | 16 Target Layouts with 1 Effector, 2 Effectors, or 4 Effectors | 6C-E |
| T5 | 11.05.2018 (810) | Single Effector Radial Target Layouts with Different Numbers of Targets | 6B |
| T5 | 09.19.2018 (763) | Closed-loop discrete decoding with a four-effector layout | 6F, 6G |
| T5 | 09.26.2018 (770) | Closed-loop discrete decoding with a two-effector layout | 6F |
| T5 | 10.01.2018 (775) | Closed-loop discrete decoding with a one-effector layout and 32-target layout | 6F, SFig 8 |
| T5 | 10.03.2018 (777) | Closed-loop discrete decoding with a four-effector layout and one-effector layout | 6F |
| T5 | 10.08.2018 (782) | Closed-loop discrete decoding with a two-effector layout and 32-target layout | 6F, SFig 8 |
| T5 | 10.10.2018 (784) | Closed-loop discrete decoding with a four-effector layout and one-effector layout | 6F |
| T5 | 10.15.2018 (789) | Closed-loop discrete decoding with a two-effector layout and 32-target layout | 6F, SFig 8 |

**Supplemental Figure 1.**
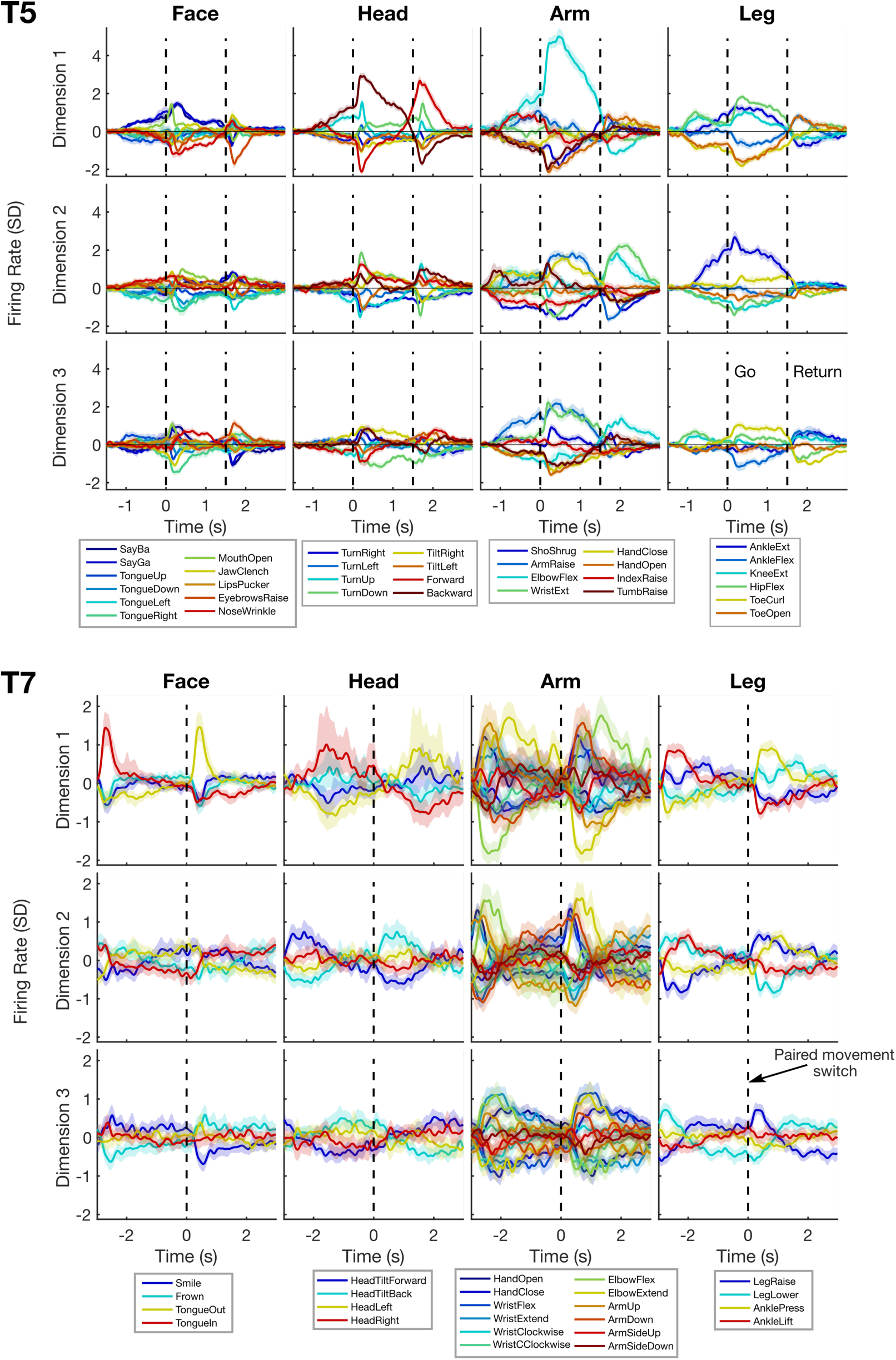
The neural population activity observed for each category of movement is rich and multi-dimensional; it cannot be explained by a generic signal that occurs for every movement. Each panel plots neural activity in one neural dimension (using the data shown in Figure 1). Each line in a panel shows the mean response to a given movement cue and shaded regions depict 95% confidence intervals. Within each column (movement set) only the neural responses for the movement cues within that set are shown (for ease of visualization). Results show that there is distinguishable neural activity (non-overlapping confidence intervals) between movements for all four movement sets and both participants. This holds true for all three neural dimensions, indicating that the movement-specific activity is strongly multi-dimensional. Note that, unlike T5, T7 performed a set of periodic paired movements (e.g. open hand, close hand, open hand, etc.). The vertical dashed lines for T7 show the paired movement “switch” time.

To make this visualization, we did a PCA-type analysis within each movement category using the data shown in Figure 1. First, the mean response of each electrode across all movements within a particular category was subtracted off. This eliminates any generic signal that varies across time but is otherwise the same across all movements, making movement-specific tuning easier to see. Next, we averaged the mean-subtracted neural activity across trials to produce a trial-averaged, three-dimensional firing rate data tensor X_NxTxM_ where N is the number of electrodes, T is the number of time bins, and M is the number of movements within a movement category. We then “unrolled” the last two dimensions of this tensor to yield a two-dimensional matrix X_NxTM_; PCA was then applied to this matrix to find the top 3 neural dimensions (principal components) that explained the most variance. The mean-subtracted neural activity was then projected into these three components and plotted above.

To generate confidence intervals, we used bootstrap resampling (percentile method, 200 re-samplings). One potential issue with applying resampling techniques to PCA is that, if multiple dimensions contain similar amounts of variance, the principle components can change order, rotate, or flip their signs from resampling to resampling, causing the confidence intervals to be much wider than they should be. To counteract this, we applied a Procrustes analysis to solve for a small 5×5 rotation matrix to align the first five principal components to a reference set computed on all the data. This problem (and solution method) is discussed in general in (Milan and Whittaker 1995).

**Supplemental Figure 2.**
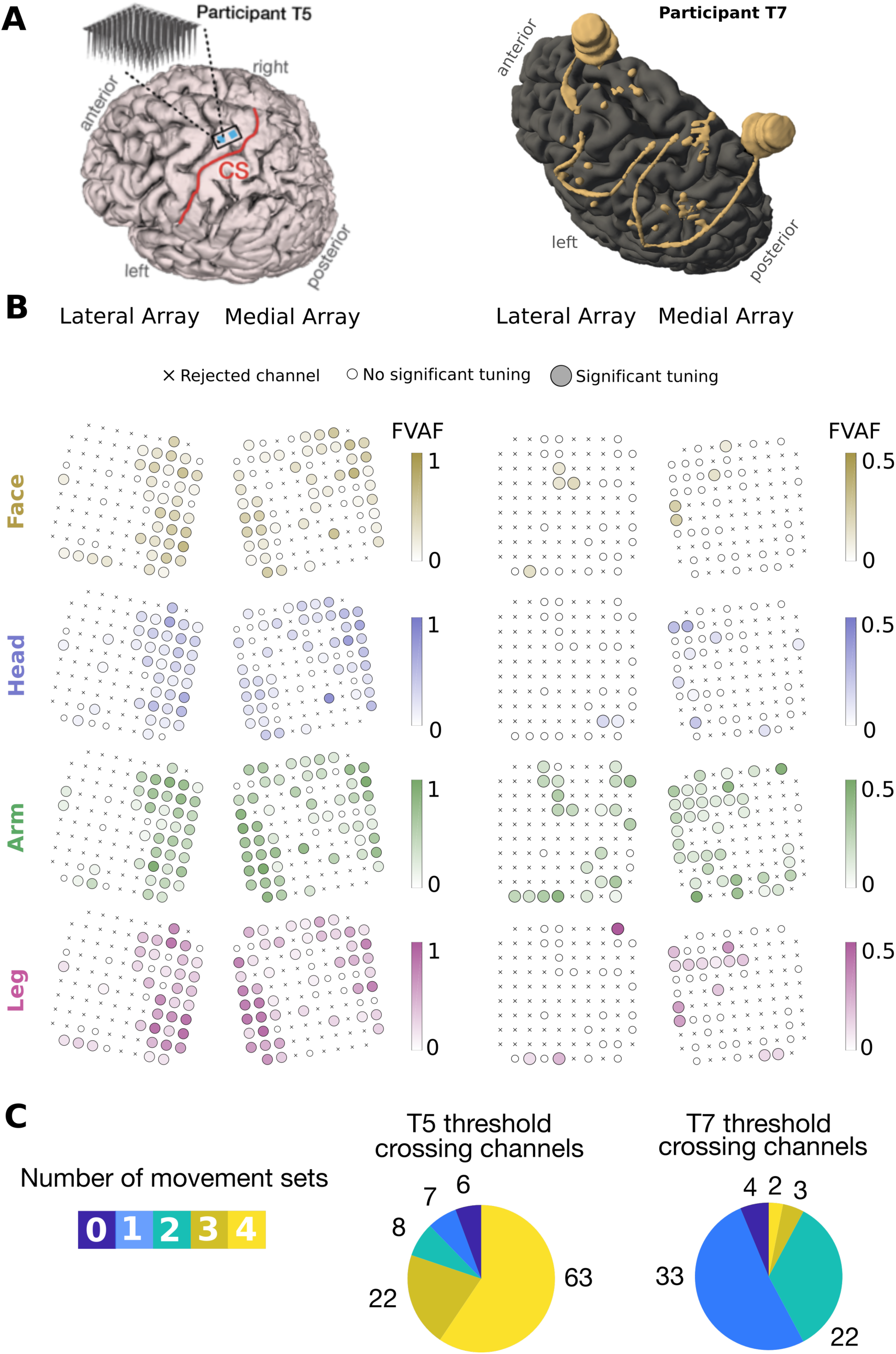
Tuning to face, head, arm and leg movements is intermixed within electrodes and appears to follow no obvious somatotopic pattern across the arrays. (A) Participants’ MRI-derived brain anatomy and microelectrode array locations. Microelectrode array locations were determined by co-registration of postoperative CT images with preoperative MRI images. For T7, the microelectrode arrays, wire bundles, and pedestal reconstructions are shown in gold. (B) The strength of each electrodes’ tuning to each category of movements is indicated with a shaded color (darker colors indicate more tuning). Tuning strength was quantified by computing the fraction of total firing rate variance accounted for by changes in firing rate due to the movement conditions. Crosses indicate “non-functioning” electrodes (mean firing rate < 1 Hz). Small white circles indicate channels that had no significant tuning to that movement category. Broad spatial tuning to all movement categories can be seen across all arrays. Additionally, tuning strength to particular movement categories does not appear to have clear gradients matching the expected somatotopy (where face, head, arm and leg tuning should proceed laterally to medially in that order), with the possible exception of leg tuning in T7. (C) Pie charts summarize the number of electrodes that had statistically significant tuning to each possible number of movement sets (from 0 to 4). In both participants, many electrodes are tuned to more than one movement set. Most of T5’s electrodes were significantly tuned to all four sets of movements, whereas most of T7’s electrodes had significant modulation to one or two sets of movements; this might be due in part to the fact that more movements were tested in T5 (especially non-arm movements) and more trials were collected (30 vs. 18 per movement).

The array maps in (B) were created using the two datasets shown in Figure 1 (10.22.2018 and 08.23.2013). To assess statistical significance and compute FVAF scores, firing rates were first computed for each trial within a 200 to 600 ms window for T5 and 200 to 1600 ms window for T7 (matching Figure 1). Next, we grouped trials into four movement sets (Head, Face, Arm and Leg) as in Figure 1. To assess the statistical significance of tuning to a movement set on a given electrode, we performed a 1-way ANOVA on all trials within that movement set. Each individual movement cue within the set was considered its own group. If the p-value was less than 0.001, the electrode was considered significantly tuned.

To compute FVAF scores for each electrode and movement set, we used the following equations:

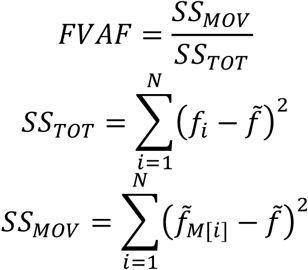

Here, SS_TOT_ is the total variance (sum of squares), SS_MOV_ is the movement-related variance, N is the total number of trials across all movements within the set, *f*_*i*_ is the firing rate for trial *i*, 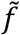 is the average firing rate across all movements within the set, and 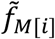 is the average firing rate for the particular movement cued on trial *i*. FVAF values range from 0 (no movement-related variance) to 1 (all variance is movement-related). T7’s FVAF shading range was reduced to 0.5 to increase visibility of the colors (since the maximum FVAF value for T7’s dataset was 0.52).

To make the pie charts in (C), the p-values from the ANOVA analysis were used to compute how many movement categories each functioning electrode was tuned to (i.e. by counting the number of p-values less than 0.001).

**Supplemental Figure 3.**
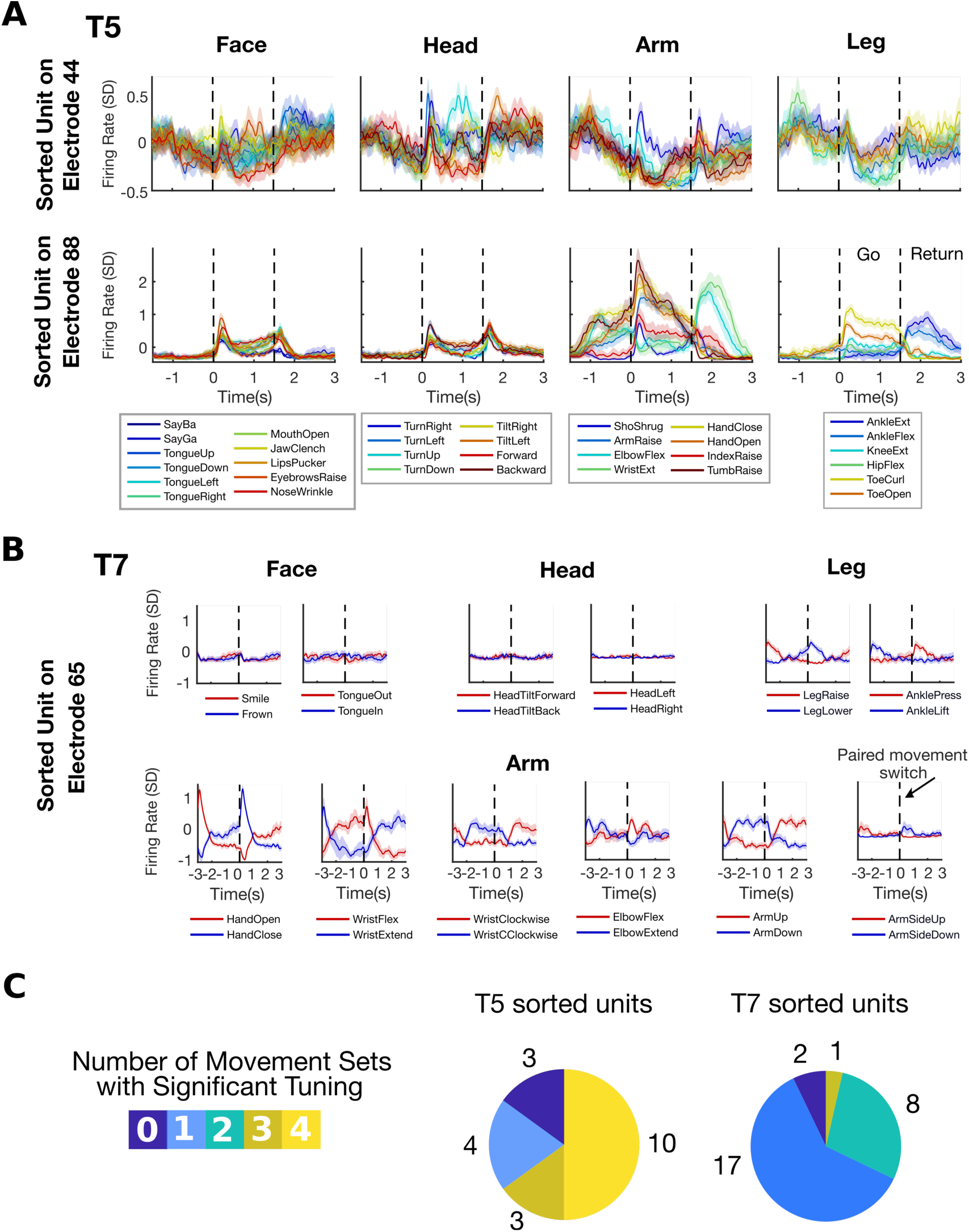
Sorted single units are tuned to multiple movement categories. (A) PSTHs for two example sorted units from participant T5. Lines show the mean firing rate and shaded regions indicate 95% CIs. The responses show broad tuning to a variety of movements across the body. (B) PSTHs for an example sorted unit from T7. Unlike T5, T7 performed a set of periodic paired movements (e.g. open hand, close hand, open hand, etc.). The vertical dashed lines show the paired movement “switch” time. Since these data were collected in a block-wise fashion (with each block testing a different movement pair), we plotted each pair separately for ease of visualization. This particular unit is strongly tuned to both the arm and the leg. (C) Pie charts summarize the number of single units that had statistically significant tuning to each possible number of movement sets (from 0 to 4). Most of T5’s sorted units modulated significantly to all four sets of movements, whereas most of T7’s sorted units had significant modulation to one or two sets of movements. This difference might be due in part to the fact that more movements were tested in T5 (especially non-arm movements) and more trials were collected (30 vs. 18 per movement).

The datasets analyzed in this figure are the same as those analyzed in Figure 1. The firing rates were smoothed with a Gaussian kernel (30 ms std for T5 and 60 ms std for T7). The statistical significance of movement tuning was assessed with a 1-way ANOVA with a p-value threshold of 0.001 (using the methods detailed in Supplemental Figure 1).

**Supplemental Figure 4.**
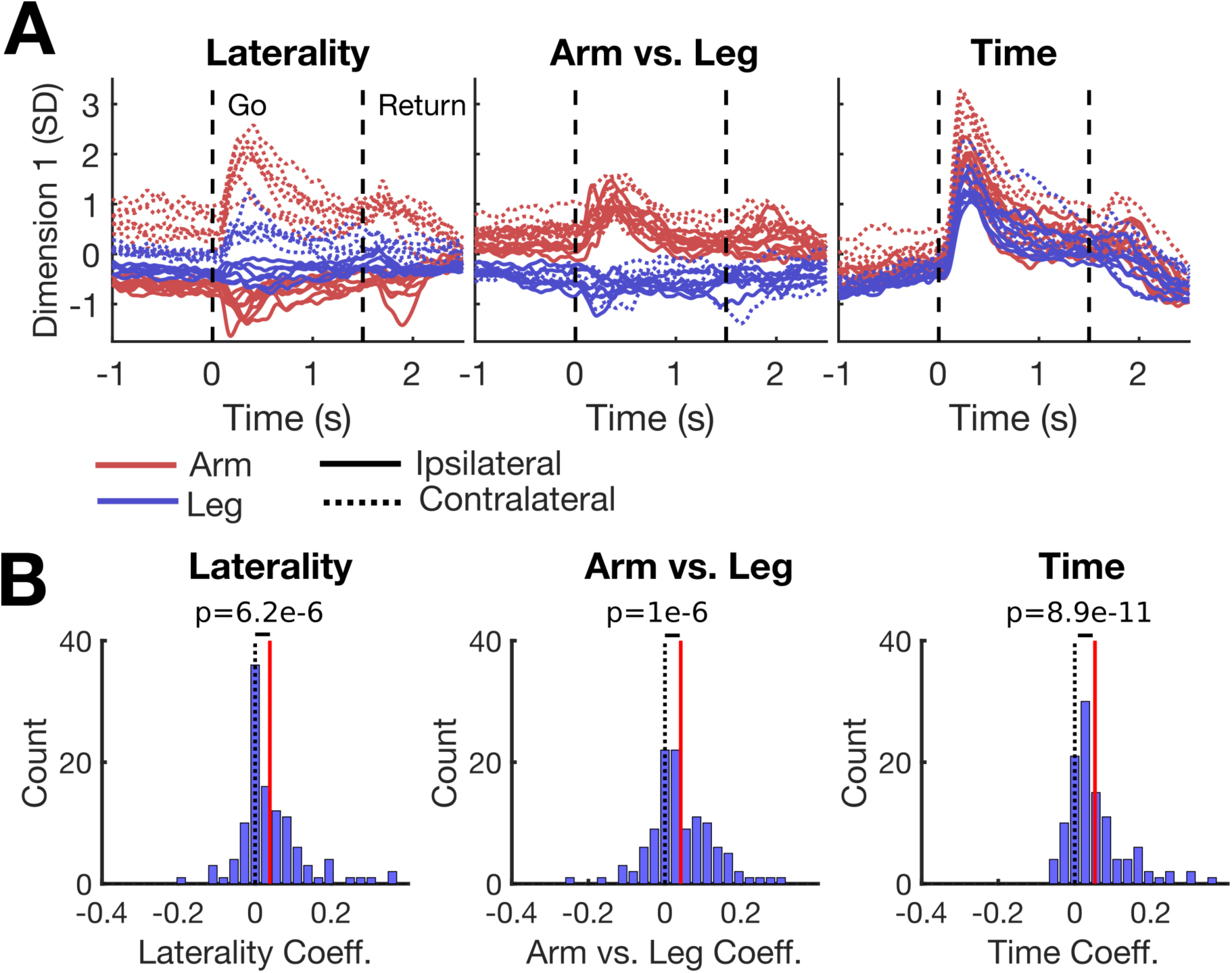
Neural dimensions coding for laterality, arm vs. leg, and timing are separable from each other (although they all describe global increases and decreases in firing rate). (A) The neural dimensions that code for laterality, arm vs. leg, and time have distinct modulation patterns and time courses. Each panel shows neural activity in a single neural dimension found by demixed PCA (dPCA), and each line shows neural activity in that dimension for a single movement condition. When finding the laterality dimension, the interaction between arm vs. leg and laterality was included along with the main effect of laterality (see below for details). The interaction can be seen mainly in the fact that the change in modulation is smaller for leg movements than arm movements. (B) The distributions of coefficients for each neural dimension are shown with histograms. The means of these distributions are all statistically significantly greater than 0 (assessed with a t-test; means are indicated with red lines and zero is indicated with black dashed lines). This implies that activity in these neural dimensions describe, in part, global increases and decreases in firing rate across the population.

The results in (B) imply that these three dimensions are all correlated with each other (i.e. they are not orthogonal). To assess alignment between the dimensions, we computed the dot product between the encoder axes associated with each of these dimensions. The dot product was 0.58 between time and laterality, 0.52 between time and arm vs. leg, and 0.54 between laterality and arm vs. leg (where 0 would mean orthogonality and 1 would mean perfect correlation).

To find these dimensions, demixed PCA was applied to the dataset shown in Fig 2 & 3 (12.05.2018) [see the Methods section on Figure 2 for more detail on dPCA]. Binned firing rates were first put into a six-dimensional data tensor with dimensions N x T x M x L x A x K, where N is the number of electrodes, T is the number of time steps, M is the number of movement conditions, L is the number of body sides (2), A signifies whether an arm or leg was moving and is equal to 2, and K is the number of trials. Note that some entries in the tensor are missing since arm and leg movements were not matched; for example, for an elbow flexion movement there are entries whenever A=1 (arm) but not A=2 (leg). When computing the marginalizations, we averaged over all available entries and skipped missing entries; this still results in a complete and exact decomposition. We grouped the factors and their interactions into the following sets for marginalizing:

Laterality Marginalization: {L, LA, LT, LAT}

Arm vs. Leg Marginalization: {A, AT}

Time Marginalization: {T}

Movement Marginalization: {M, ML, MA, MAL, MT, MLT, MAT, MALT}

**Supplemental Figure 5.**
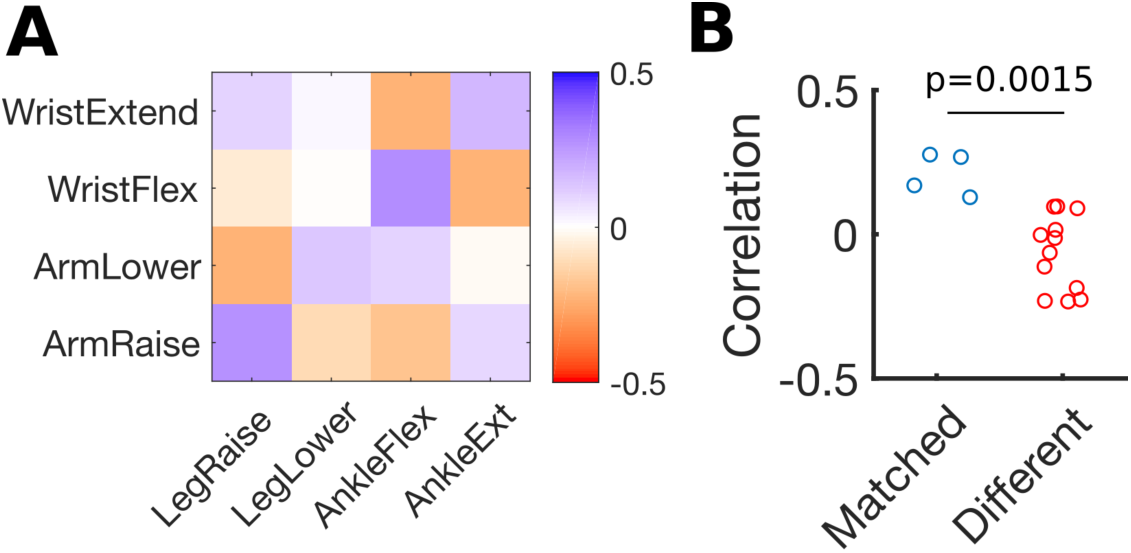
Homologous arm and leg movements are correlated for participant T7. (A) The correlations between firing rate vectors associated with each leg and arm movement are indicated in color (same methods as in Fig 3). Only four leg movements were tested with T7. The diagonal stripe of blue indicates that homologous movements are positively correlated. A 0 to 1000 ms time window aligned to the paired movement switching cue was used to compute the firing rate vectors for the correlations. (B) The distributions of correlation coefficients are plotted for matching movements (the diagonal of the correlation matrix) and different movements (all other values). Jitter was added to aid visualization. A t-test indicates statistical significance between the two distributions (r=0.22 vs. -0.07).

**Supplemental Figure 6.**
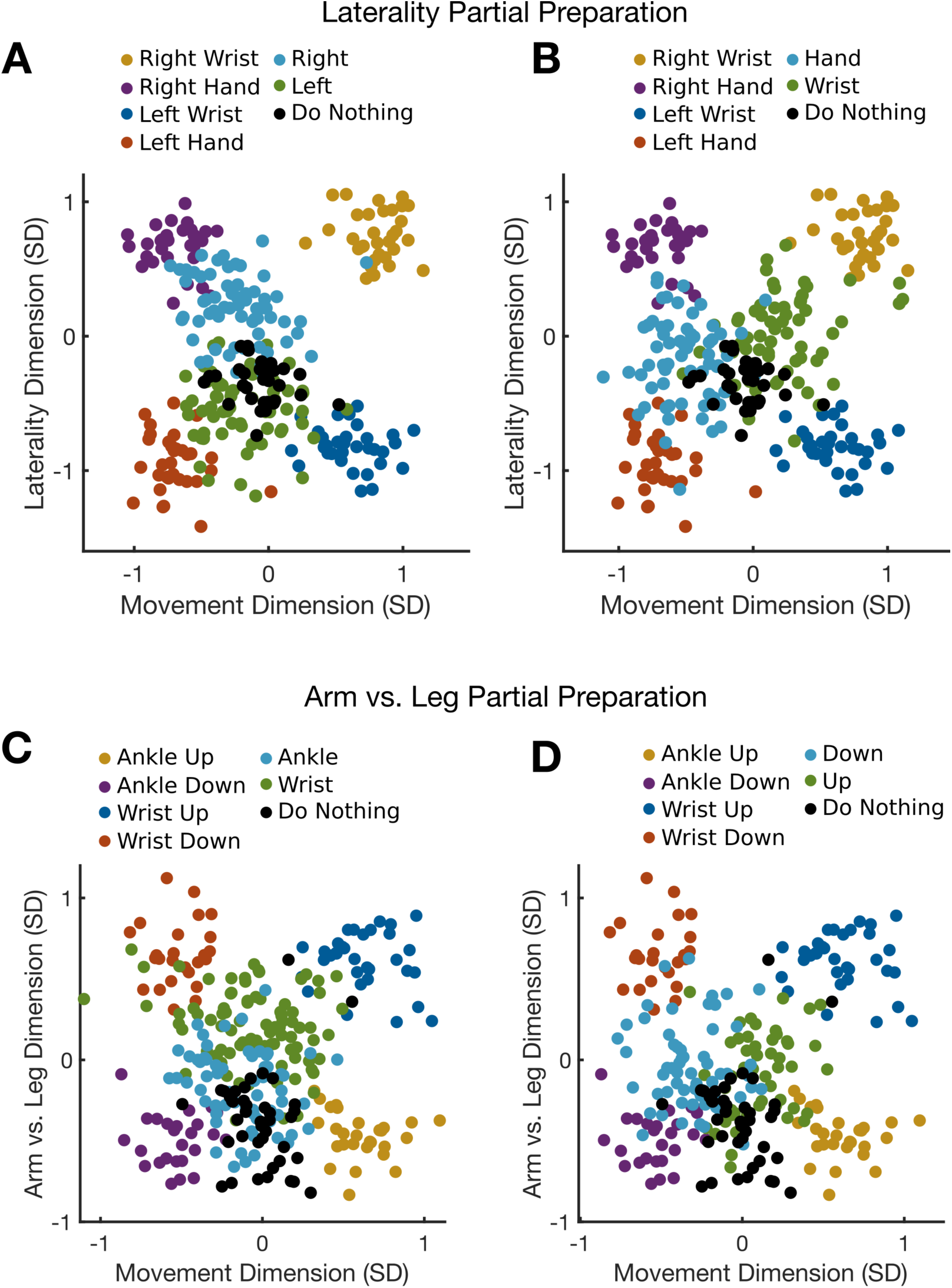
Single trial distributions of neural activity during the instructed delay period of the partially cued movement task show patterns consistent with genuine partial preparation as opposed to a mixture of full preparation and no preparation. (A, B) Single trial distributions of neural activity for the partially cued laterality and movement task; each panel depicts different partially cued distributions for ease of visualization. Each dot shows the neural activity in the laterality and movement dimensions in a 300 ms window just prior to the go cue. Each color corresponds to a different cue. If the partial preparation we observed during the partially-cued trials were an artifact of T5 deciding sometimes to prepare one of the two potential upcoming movements (e.g. preparing either the right wrist or the right hand movement once the right side of the body was cued), and other times to prepare nothing, we would expect to see dots from the “right” partial cue condition (light blue dots in panel A) to occupy either the right hand distribution (purple dots), right wrist distribution (gold dots), or the “do nothing” distribution (black dots). Instead, the distribution appears to be in a genuinely different and “in-between” location that cannot be explained by a Gaussian mixture distribution composed of the fully-specified and “do nothing” distributions. (C, D) Single trial distributions of neural activity for the partially cued arm vs. leg and movement task, depicted as in (A, B). Similar to the distributions for the laterality task, the partially-cued distributions appear to lie in “in-between” locations that cannot be explained by sometimes fully preparing possible upcoming movements and sometimes preparing nothing.

**Supplemental Figure 7.**
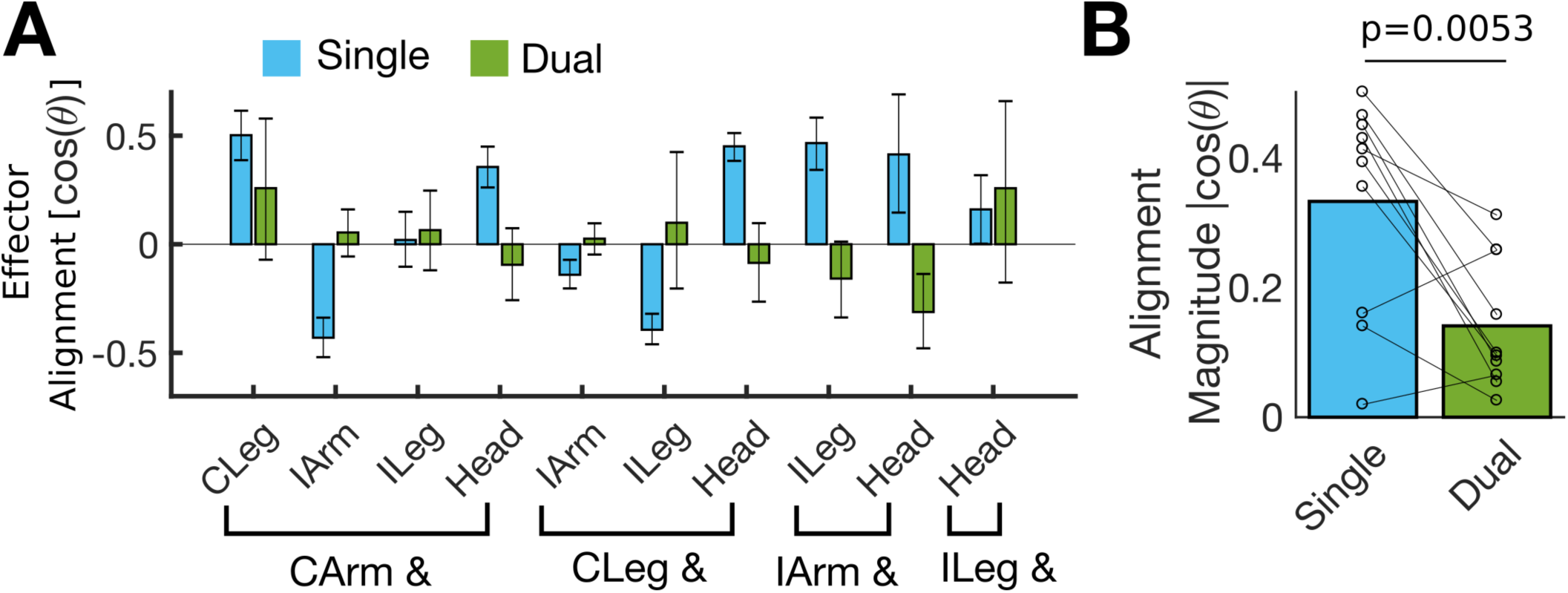
The axes of neural representation during dual effector movement change so that the effectors become less aligned with each other. (A) Alignment between the neural axes of both effectors is shown for the single movement trials (blue bars) and the simultaneous movement trials (green bars). Alignment was measured by computing the cosine of the angle between the two vectors (same metric used as in Fig 5F, bottom panel). Overall, alignment appears to move closer to zero in the dual movement case, meaning that the axes of representation have become more decorrelated. Note that significant positive alignments can be seen for cases of matching movements of homologous joints (e.g. CArm and CLeg) and negative alignments can be seen for cases of opposite movements of homologous joints (e.g. CArm and IArm), consistent with what was seen in Fig. 4. (B) The alignment magnitude (absolute value of the cosine of the angle) is summarized for the single movement context and the dual movement context; each circle shows data from a single pairing and lines connect matching pairs. Bar heights indicate the mean across all pairings. A paired t-test reveals a statistically significant difference in alignment between the single and dual contexts (p=0.013; 0.27 vs. 0.12).

**Supplemental Figure 8.**
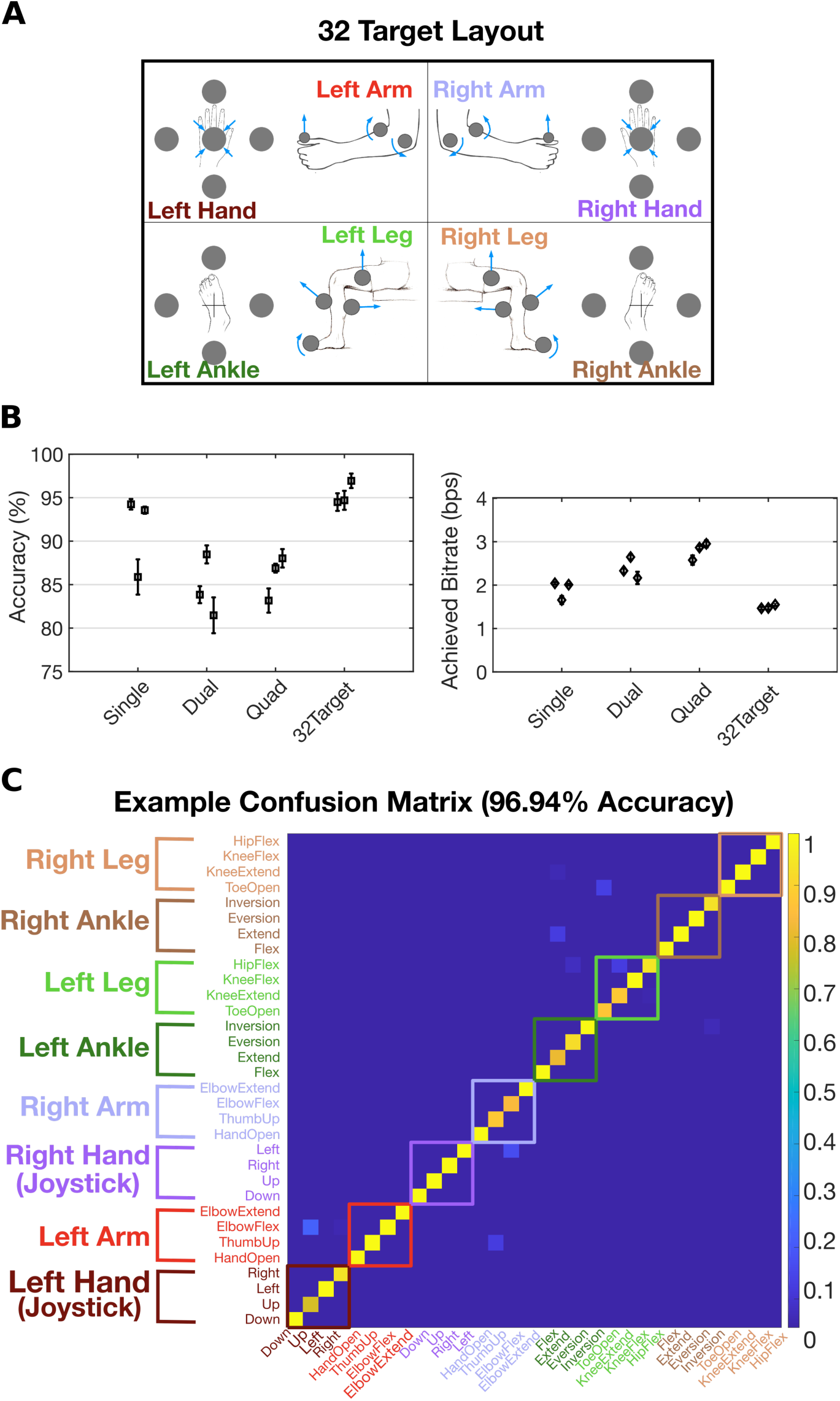
Closed-loop discrete decoding with a 32-target layout. The decoding methods used for the 32-target layout were the same as those used in Figure 6, except that an additional two second instructed delay period was added before each trial to inform T5 of the upcoming target. This allowed T5 enough time to recognize and prepare the movement (Supplemental Video 2 shows example trials). During this time, the target appeared red. After the delay period, the target turned blue and T5 had one second to make the movement. The discrete decoder used data only from this one second movement period to classify the movement. We found that, without the delay period, T5 could not recognize the movement quickly enough and perform it correctly within the allotted time.

(A) In addition to the pointing movements at the wrists and ankles from the quad-effector 16 target layout, the following additional 16 targets were added: hand grasp, elbow flexion & extension, thumb raise, toe spread, knee flexion & extension, and thigh raise.

(B) Accuracy and achieved bit rate for the 32-target configuration (with the single, dual, and quad-effector performance data reproduced from Figure 6 for reference). Accuracy is high (mean of 95% across all three sessions) but the achieved bit rate is relatively low (mean of 1.5 bps) because of the additional two second delay period. Without the two second delay period, the bit rate would have been 4.52 bps, suggesting that high bit rates are possible if the user can learn to select between a large number of movements quickly.

(C) An example confusion matrix summarizing the online decoding performance for session 10.15.2018. Each entry (i, j) in the matrix is colored according to the fraction of trials where movement j was decoded (out of all trials where movement i was cued).

**Supplemental Figure 9.**
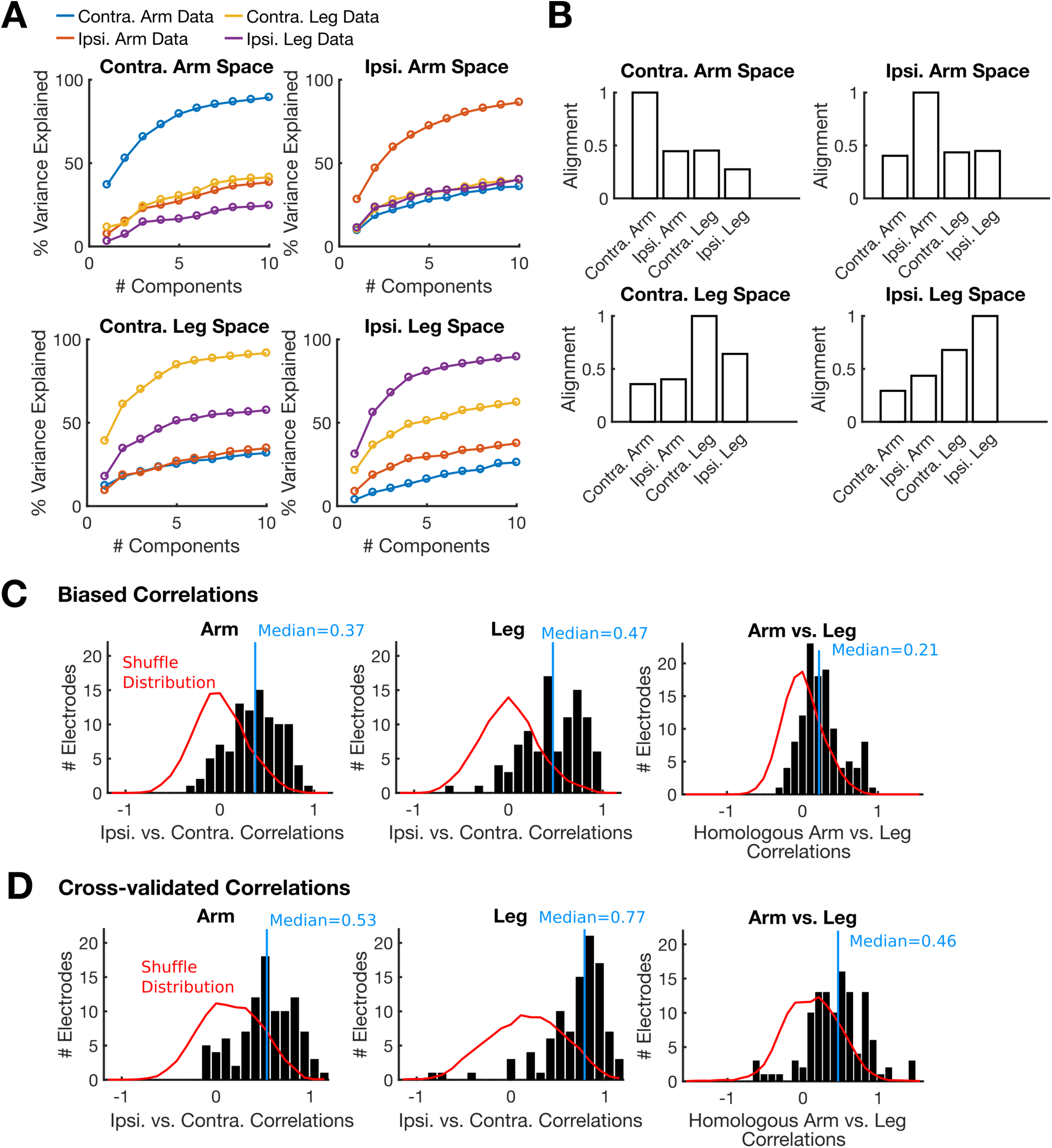
A comparison of ipsilateral vs. contralateral correlation strength to recent studies of macaque M1 shows that the correlations observed in this study are stronger. We computed the same correlation metrics used in two recent studies (1,2) in order to make a direct comparison of correlation strengths [compare this figure to Figure 7 E-G, L-N in (1) and Figure 4C in (2)]. Results show that the strength of ipsilateral vs. contralateral (and arm vs. leg) neural correlations are stronger here than what was observed in (1,2), consistent with human precentral gyrus being homologous to macaque PMd rather than M1. See (1,2) for more detailed explanations of how each metric was computed.

(A) This plot replicates Figure 7 E-F and L-M from (1). Each panel shows the % variance explained of the trial-averaged neural data (within a 200 to 1000 ms time window after the go cue) as a function of the number of principal components used. The principal components (PCs) for each panel were derived from a different set of movements (contra. arm, ipsi. arm, contra. leg, or ipsi. leg) and then applied to all other sets of movements (using the same dataset analyzed in Figure 2 of this study). Results show that each set of PCs can explain considerable amounts of variance related to other movement types, more so than what was found in (1); nevertheless, the spaces are still clearly distinct from each other.

(B) This plot replicates Figure 7G and 7N from (1). Each panel summarizes the “alignment” of one subspace (as defined by its first 10 principal components) with each other subspace [see (1) for methods details]. The alignment score is equal to the ratio of the variance explained of dataset A in subspace S divided by the total possible explainable variance using the top 10 PCs of A’s own subspace. Subspace alignment is 0.42 on average for the arms, 0.65 for the legs, and 0.41 for arms vs. legs on the same body side. In (1), subspace alignment between the arms was 0.19 for monkey P and 0.29 for monkey M. Since variance explained is a “squared” metric, an alignment score of 0.42 implies that, on average, a neural signal of size 1 would shrink only to size 0.65 when projected into the opposite arm’s subspace [since sqrt(0.42)=0.65].

(C) This plot replicates Figure 4C from (2) and summarizes the distribution of ipsilateral vs. contralateral (and arm vs. leg) correlations for each electrode. Data was smoothed as in (2) [using a 25 ms standard deviation Gaussian kernel]. The trial-averaged responses for each movement were concatenated together, mean-subtracted, and then correlated across ipsilateral and contralateral sets of movements. A 200 to 1000 ms window after the go cue was used for the analysis. The correlations appear consistently more positive than in Figure 4C from (2) [where the median correlations were reported as 0.16 and 0.08 for monkey E and F]. Since these correlations were computed in a “standard” way (where the trial-averaged responses are directly correlated together with no cross-validation employed), they are labeled as “biased”, since this method systematically biases the correlations downwards (because sampling error reduces the correlation).

(D) We computed the correlations again using the cross-validated approach we described in the Methods. Interestingly, the correlations substantially increase, suggesting that single-electrode or single-neuron responses may not always have a high enough signal-to-noise ratio to enable accurate estimation of correlations using the standard approach.

Note that all distributions of correlation coefficients in (C) and (D) were highly statistically significant (p<1e-13 in all cases; a sign test was used to test whether the median was different from 0).

(1) Heming, Ethan A., Kevin P. Cross, Tomohiko Takei, Douglas J. Cook, and Stephen H. Scott. “Keep Your Hands Apart: Independent Representations of Ipsilateral and Contralateral Forelimbs in Primary Motor Cortex.” BioRxiv, March 24, 2019, 587378. https://doi.org/10.1101/587378.

(2) Ames, Katherine Cora, and Mark M. Churchland. “Motor Cortex Signals Corresponding to the Two Arms Are Shared across Hemispheres, Mixed among Neurons, yet Partitioned within the Population Response.” BioRxiv, February 18, 2019, 552257. https://doi.org/10.1101/552257.

**Supplemental Figure 10.**
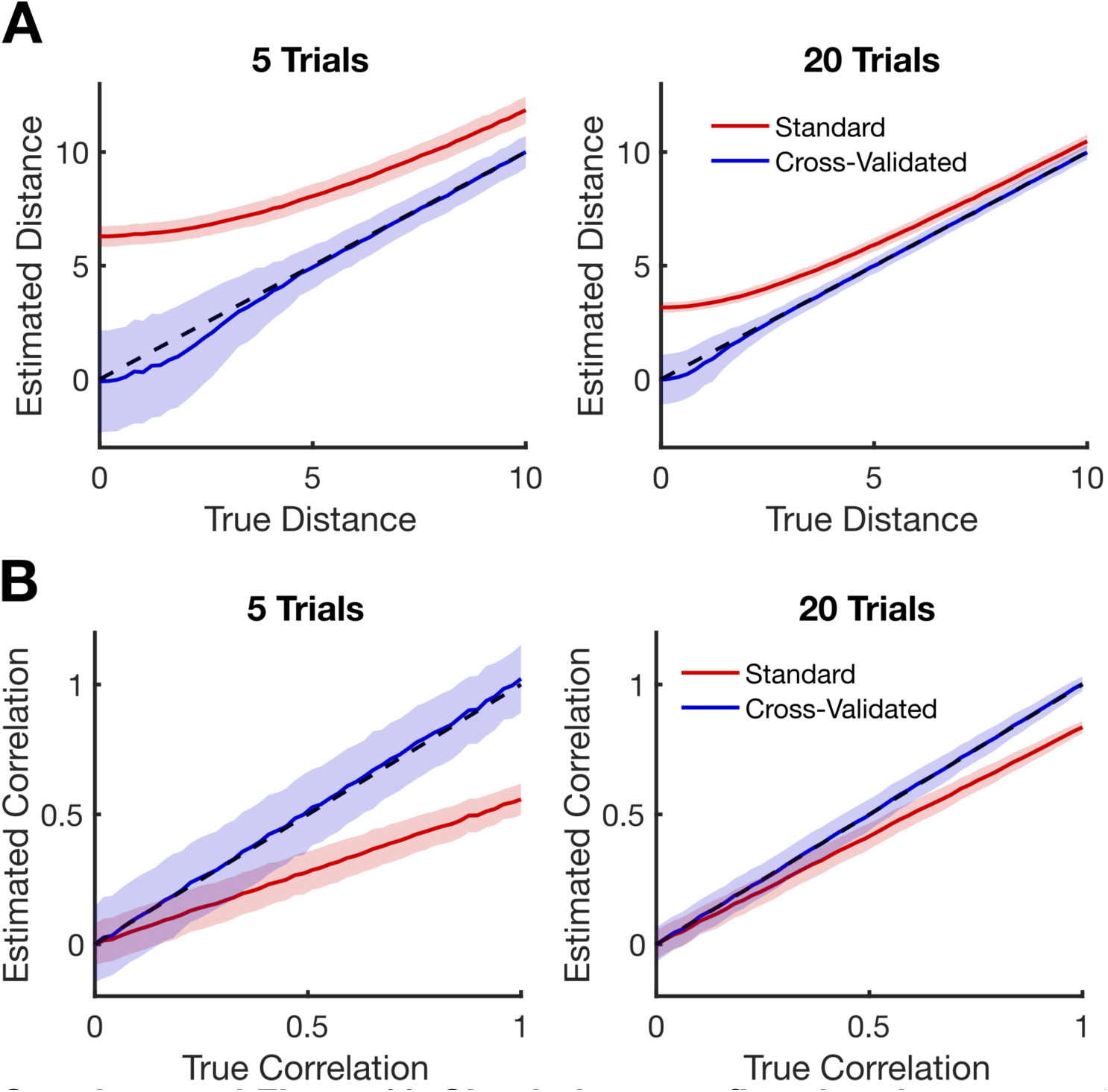
Simulations confirm that the cross-validated metrics used in this study substantially reduce bias. (A) An assessment of how the cross-validated distance estimation approach used here performs compared to a standard approach when estimating the distance between two multivariate distributions of firing rate vectors (e.g. as computed in Fig 1C). Lines show the mean of each estimator across 1,000 repetitions and shaded regions show ± 1 standard deviations. Dashed black lines show the identity line. The cross-validated metric lies close to the identity line, although its variance is larger for small distances. For very small distances, the metric is conservative (it lies slightly below the identity line). In contrast, the standard method is heavily biased upwards because sampling error in the sample means causes the distance metric to become inflated. As the signal-to-noise ratio of the data increases (in the case of more samples or larger distances), the difference between the approaches becomes smaller. The cross-validated metric substantially extends the range over which accurate results can be obtained and provides assurance that non-zero distances, if statistically significant, likely reflect a true difference in the distributions.

Simulated firing rates were generated for distribution A by sampling from a multivariate normal distribution *N*(**0, I**_***n***_), where **I**_***n***_ is the identity matrix (with n=100 dimensions). Neural data from distribution B was sampled from 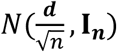, where ***d*** is a vector containing the scalar value *d* in all entries (which causes distribution B to be a distance of *d* from distribution A). After generating data, the distance between the A and B distributions was estimated either in a “standard” way by computing the Euclidean norm of the difference between their sample means, or in a cross-validated median-unbiased way (as described in the Methods). We repeated the simulation 1,000 times for each of 50 distances, with either 5 samples from each distribution (left panel) or 20 samples (right panel).

(B) An assessment of how the cross-validated distance estimation approach used here performs compared to a standard approach when estimating the correlation between the means of two distributions of firing rate vectors (e.g. as computed in Fig 3B-C). Lines show the mean of each estimator across 1,000 repetitions and shaded regions show ± 1 standard deviations. Dashed black lines show the identity line. The cross-validated metric lies much closer to the identity line; in contrast, the standard method is biased downwards since sampling error causes the sample means to be less correlated than the true means really are. For a more applied example of how cross-validated correlations can sometimes make a substantial difference, see Supplemental Figure XC-D.

To generate these simulated results, the performance of each approach was assessed on 50 different correlation values ranging from 0 to 1. To generate simulated data, we first randomly generated the 100-dimensional mean vectors of each distribution so that they had the desired correlation and a Euclidean norm of 5 (but were otherwise random). To do so, the 100-dimensional mean vector for distribution A was first generated by sampling each coefficient from a normal distribution, subtracting the mean across the whole vector, and then dividing by the vector norm to produce a centered vector with a norm of 1. We then repeated this same process again, yielding another random vector R with norm 1. We then set the mean of distribution B equal to a weighted sum of the mean of A and R so that the resulting vector also had a norm of 5 and was correlated with the mean of A the desired amount.

Simulated firing rates were then sampled from distributions A and B (each distribution had a diagonal covariance matrix equal to the identity matrix). After generating data, the correlation between the A and B distributions was estimated either in a “standard” way by computing the correlation between their sample means, or in a cross-validated way (as described in the Methods). We repeated the simulation 1,000 times for each of 50 correlation values, with either 5 samples from each class (left panel) or 20 samples (right panel).

## References

Aflalo, Tyson, Spencer Kellis, Christian Klaes, Brian Lee, Ying Shi, Kelsie Pejsa, Kathleen Shanfield, et al. 2015. “Decoding Motor Imagery from the Posterior Parietal Cortex of a Tetraplegic Human.” Science 348 (6237): 906–10. https://doi.org/10.1126/science.aaa5417.

Ajiboye, A Bolu, Francis R Willett, Daniel R Young, William D Memberg, Brian A Murphy, Jonathan P Miller, Benjamin L Walter, et al. 2017. “Restoration of Reaching and Grasping Movements through Brain-Controlled Muscle Stimulation in a Person with Tetraplegia: A Proof-of-Concept Demonstration.” The Lancet 389 (10081): 1821–30. https://doi.org/10.1016/S0140-6736(17)30601-3.

Allefeld, Carsten, and John-Dylan Haynes. 2014. “Searchlight-Based Multi-Voxel Pattern Analysis of FMRI by Cross-Validated MANOVA.” NeuroImage 89 (April): 345–57. https://doi.org/10.1016/j.neuroimage.2013.11.043.

Ames, Katherine Cora, and Mark M. Churchland. 2019. “Motor Cortex Signals Corresponding to the Two Arms Are Shared across Hemispheres, Mixed among Neurons, yet Partitioned within the Population Response.” BioRxiv, February, 552257. https://doi.org/10.1101/552257.

Bouton, Chad E., Ammar Shaikhouni, Nicholas V. Annetta, Marcia A. Bockbrader, David A. Friedenberg, Dylan M. Nielson, Gaurav Sharma, et al. 2016. “Restoring Cortical Control of Functional Movement in a Human with Quadriplegia.” Nature 533 (7602): 247–50. https://doi.org/10.1038/nature17435.

Bundy, David T., Nicholas Szrama, Mrinal Pahwa, and Eric C. Leuthardt. 2018. “Unilateral, 3D Arm Movement Kinematics Are Encoded in Ipsilateral Human Cortex.” Journal of Neuroscience 38 (47): 10042–56. https://doi.org/10.1523/JNEUROSCI.0015-18.2018.

Christou, Evangelos A., and Tiffany M. Rodriguez. 2008. “Time but Not Force Is Transferred Between Ipsilateral Upper and Lower Limbs.” Journal of Motor Behavior 40 (3): 186–89. https://doi.org/10.3200/JMBR.40.3.186-189.

Cisek, Paul, Donald J. Crammond, and John F. Kalaska. 2003. “Neural Activity in Primary Motor and Dorsal Premotor Cortex In Reaching Tasks With the Contralateral Versus Ipsilateral Arm.” Journal of Neurophysiology 89 (2): 922–42. https://doi.org/10.1152/jn.00607.2002.

Collinger, Jennifer L, Brian Wodlinger, John E Downey, Wei Wang, Elizabeth C Tyler-Kabara, Douglas J Weber, Angus JC McMorland, Meel Velliste, Michael L Boninger, and Andrew B Schwartz. 2013. “High-Performance Neuroprosthetic Control by an Individual with Tetraplegia.” The Lancet 381 (9866): 557–64. https://doi.org/10.1016/S0140-6736(12)61816-9.

Criscimagna-Hemminger, Sarah E., Opher Donchin, Michael S. Gazzaniga, and Reza Shadmehr. 2003. “Learned Dynamics of Reaching Movements Generalize From Dominant to Nondominant Arm.” Journal of Neurophysiology 89 (1): 168–76. https://doi.org/10.1152/jn.00622.2002.

Crone, N. E., D. L. Miglioretti, B. Gordon, and R. P. Lesser. 1998. “Functional Mapping of Human Sensorimotor Cortex with Electrocorticographic Spectral Analysis. II. Event-Related Synchronization in the Gamma Band.” Brain 121 (12): 2301–15. https://doi.org/10.1093/brain/121.12.2301.

Crone, N. E., D. L. Miglioretti, B. Gordon, J. M. Sieracki, M. T. Wilson, S. Uematsu, and R. P. Lesser. 1998. “Functional Mapping of Human Sensorimotor Cortex with Electrocorticographic Spectral Analysis. I. Alpha and Beta Event-Related Desynchronization.” Brain 121 (12): 2271–99. https://doi.org/10.1093/brain/121.12.2271.

Diedrichsen, Jörn, Tobias Wiestler, and John W. Krakauer. 2013. “Two Distinct Ipsilateral Cortical Representations for Individuated Finger Movements.” Cerebral Cortex 23 (6): 1362–77. https://doi.org/10.1093/cercor/bhs120.

Fujiwara, Yusuke, Riki Matsumoto, Takuro Nakae, Kiyohide Usami, Masao Matsuhashi, Takayuki Kikuchi, Kazumichi Yoshida, et al. 2017. “Neural Pattern Similarity between Contra- and Ipsilateral Movements in High-Frequency Band of Human Electrocorticograms.” NeuroImage 147 (February): 302–13. https://doi.org/10.1016/j.neuroimage.2016.11.058.

Ganguly, Karunesh, Lavi Secundo, Gireeja Ranade, Amy Orsborn, Edward F. Chang, Dragan F. Dimitrov, Jonathan D. Wallis, Nicholas M. Barbaro, Robert T. Knight, and Jose M. Carmena. 2009. “Cortical Representation of Ipsilateral Arm Movements in Monkey and Man.” Journal of Neuroscience 29 (41): 12948–56. https://doi.org/10.1523/JNEUROSCI.2471-09.2009.

Genon, Sarah, Hai Li, Lingzhong Fan, Veronika I. Müller, Edna C. Cieslik, Felix Hoffstaedter, Andrew T. Reid, et al. 2017. “The Right Dorsal Premotor Mosaic: Organization, Functions, and Connectivity.” Cerebral Cortex (New York, NY) 27 (3): 2095–2110. https://doi.org/10.1093/cercor/bhw065.

Genon, Sarah, Andrew Reid, Hai Li, Lingzhong Fan, Veronika I. Müller, Edna C. Cieslik, Felix Hoffstaedter, et al. 2018. “The Heterogeneity of the Left Dorsal Premotor Cortex Evidenced by Multimodal Connectivity-Based Parcellation and Functional Characterization.” NeuroImage, Segmenting the Brain, 170 (April): 400–411. https://doi.org/10.1016/j.neuroimage.2017.02.034.

Georgopoulos, A. P., J. F. Kalaska, R. Caminiti, and J. T. Massey. 1982. “On the Relations between the Direction of Two-Dimensional Arm Movements and Cell Discharge in Primate Motor Cortex.” The Journal of Neuroscience 2 (11): 1527–1537.

Geyer, S., M. Matelli, G. Luppino, and K. Zilles. 2000. “Functional Neuroanatomy of the Primate Isocortical Motor System.” Anatomy and Embryology 202 (6): 443–74.

Heming, Ethan A., Kevin P. Cross, Tomohiko Takei, Douglas J. Cook, and Stephen H. Scott. 2019. “Keep Your Hands Apart: Independent Representations of Ipsilateral and Contralateral Forelimbs in Primary Motor Cortex.” BioRxiv, March, 587378. https://doi.org/10.1101/587378.

Hochberg, Leigh R., Daniel Bacher, Beata Jarosiewicz, Nicolas Y. Masse, John D. Simeral, Joern Vogel, Sami Haddadin, et al. 2012. “Reach and Grasp by People with Tetraplegia Using a Neurally Controlled Robotic Arm.” Nature 485 (7398): 372–75. https://doi.org/10.1038/nature11076.

Ifft, Peter J, Solaiman Shokur, Zheng Li, Mikhail A Lebedev, and Miguel A L Nicolelis. 2013. “A Brain-Machine Interface Enables Bimanual Arm Movements in Monkeys.” Science Translational Medicine 5 (210): 210ra154. https://doi.org/10.1126/scitranslmed.3006159.

Jin, Y., M. Lu, X. Wang, S. Zhang, J. Zhu, and X. Zheng. 2016. “Electrocorticographic Signals Comparison in Sensorimotor Cortex between Contralateral and Ipsilateral Hand Movements.” In 2016 38th Annual International Conference of the IEEE Engineering in Medicine and Biology Society (EMBC), 1544–47. https://doi.org/10.1109/EMBC.2016.7591005.

Kelso, J. a. S., and P. G. Zanone. 2002. “Coordination Dynamics of Learning and Transfer across Different Effector Systems.” Journal of Experimental Psychology. Human Perception and Performance 28 (4): 776–97.

Kobak, Dmitry, Wieland Brendel, Christos Constantinidis, Claudia E. Feierstein, Adam Kepecs, Zachary F. Mainen, Xue-Lian Qi, Ranulfo Romo, Naoshige Uchida, and Christian K. Machens. 2016. “Demixed Principal Component Analysis of Neural Population Data.” ELife 5 (April): e10989. https://doi.org/10.7554/eLife.10989.

Kurata, K. 1989. “Distribution of Neurons with Set- and Movement-Related Activity before Hand and Foot Movements in the Premotor Cortex of Rhesus Monkeys.” Experimental Brain Research 77 (2): 245–56. https://doi.org/10.1007/BF00274982.

Latash, Mark L. 1999. “Mirror Writing: Learning, Transfer, and Implications for Internal Inverse Models.” Journal of Motor Behavior 31 (2): 107–11. https://doi.org/10.1080/00222899909600981.

Leyton, A. S. F., and C. S. Sherrington. 1917. “Observations on the Excitable Cortex of the Chimpanzee, Orang-Utan, and Gorilla.” Quarterly Journal of Experimental Physiology 11 (2): 135–222. https://doi.org/10.1113/expphysiol.1917.sp000240.

Lotze, M., M. Erb, H. Flor, E. Huelsmann, B. Godde, and W. Grodd. 2000. “FMRI Evaluation of Somatotopic Representation in Human Primary Motor Cortex.” NeuroImage 11 (5): 473–81. https://doi.org/10.1006/nimg.2000.0556.

Meier, Jeffrey D., Tyson N. Aflalo, Sabine Kastner, and Michael S. A. Graziano. 2008. “Complex Organization of Human Primary Motor Cortex: A High-Resolution FMRI Study.” Journal of Neurophysiology 100 (4): 1800–1812. https://doi.org/10.1152/jn.90531.2008.

Milan, Luis, and Joe Whittaker. 1995. “Application of the Parametric Bootstrap to Models That Incorporate a Singular Value Decomposition.” Journal of the Royal Statistical Society: Series C (Applied Statistics) 44 (1): 31–49. https://doi.org/10.2307/2986193.

Miller, Kai J., Eric C. Leuthardt, Gerwin Schalk, Rajesh P. N. Rao, Nicholas R. Anderson, Daniel W. Moran, John W. Miller, and Jeffrey G. Ojemann. 2007. “Spectral Changes in Cortical Surface Potentials during Motor Movement.” Journal of Neuroscience 27 (9): 2424–32. https://doi.org/10.1523/JNEUROSCI.3886-06.2007.

Morris, Tiffany, Nicki Ann Newby, Michael Wininger, and William Craelius. 2009. “Inter-Limb Transfer of Learned Ankle Movements.” Experimental Brain Research 192 (1): 33–42. https://doi.org/10.1007/s00221-008-1547-x.

Musallam, S., B. D. Corneil, B. Greger, H. Scherberger, and R. A. Andersen. 2004. “Cognitive Control Signals for Neural Prosthetics.” Science 305 (5681): 258–62. https://doi.org/10.1126/science.1097938.

Nozaki, Daichi, Isaac Kurtzer, and Stephen H. Scott. 2006. “Limited Transfer of Learning between Unimanual and Bimanual Skills within the Same Limb.” Nature Neuroscience 9 (11): 1364–66. https://doi.org/10.1038/nn1785.

Nuyujukian, Paul, Joline M. Fan, Jonathan C. Kao, Stephen I. Ryu, and Krishna V. Shenoy. 2015. “A High-Performance Keyboard Neural Prosthesis Enabled by Task Optimization.” IEEE Transactions on Bio-Medical Engineering 62 (1): 21–29. https://doi.org/10.1109/TBME.2014.2354697.

Pandarinath, Chethan, Paul Nuyujukian, Christine H. Blabe, Brittany L. Sorice, Jad Saab, Francis R. Willett, Leigh R. Hochberg, Krishna V. Shenoy, and Jaimie M. Henderson. 2017. “High Performance Communication by People with Paralysis Using an Intracortical Brain-Computer Interface.” ELife 6 (February): e18554. https://doi.org/10.7554/eLife.18554.

Penfield, W., and E. Boldrey. 1937. “Somatic Motor and Sensory Representation in the Cerebral Cortex of Man as Studied by Electrical Stimulation.” Brain: A Journal of Neurology 60: 389–443. https://doi.org/10.1093/brain/60.4.389.

Penfield, Wilder, and Theodore Rasmussen. 1950. The Cerebral Cortex of Man; a Clinical Study of Localization of Function. The Cerebral Cortex of Man; a Clinical Study of Localization of Function. Oxford, England: Macmillan.

Qi, Hui-Xin, Iwona Stepniewska, and Jon H. Kaas. 2000. “Reorganization of Primary Motor Cortex in Adult Macaque Monkeys With Long-Standing Amputations.” Journal of Neurophysiology 84 (4): 2133–47. https://doi.org/10.1152/jn.2000.84.4.2133.

Rizzolatti, G., G. Luppino, and M. Matelli. 1998. “The Organization of the Cortical Motor System: New Concepts.” Electroencephalography and Clinical Neurophysiology 106 (4): 283–96.

Rokni, Uri, Orna Steinberg, Eilon Vaadia, and Haim Sompolinsky. 2003. “Cortical Representation of Bimanual Movements.” Journal of Neuroscience 23 (37): 11577–86. https://doi.org/10.1523/JNEUROSCI.23-37-11577.2003.

Ruescher, Johanna, Olga Iljina, Dirk-Matthias Altenmüller, Ad Aertsen, Andreas Schulze-Bonhage, and Tonio Ball. 2013. “Somatotopic Mapping of Natural Upper- and Lower-Extremity Movements and Speech Production with High Gamma Electrocorticography.” NeuroImage 81 (November): 164–77. https://doi.org/10.1016/j.neuroimage.2013.04.102.

Santhanam, Gopal, Stephen I. Ryu, Byron M. Yu, Afsheen Afshar, and Krishna V. Shenoy. 2006. “A High-Performance Brain–Computer Interface.” Nature 442 (7099): 195–98. https://doi.org/10.1038/nature04968.

Savin, Douglas N., and Susanne M. Morton. 2008. “Asymmetric Generalization between the Arm and Leg Following Prism-Induced Visuomotor Adaptation.” Experimental Brain Research 186 (1): 175–82. https://doi.org/10.1007/s00221-007-1220-9.

Schieber, M. H. 2001. “Constraints on Somatotopic Organization in the Primary Motor Cortex.” Journal of Neurophysiology 86 (5): 2125–43. https://doi.org/10.1152/jn.2001.86.5.2125.

Severiano, Ana, João A. Carriço, D. Ashley Robinson, Mário Ramirez, and Francisco R. Pinto. 2011. “Evaluation of Jackknife and Bootstrap for Defining Confidence Intervals for Pairwise Agreement Measures.” PLOS ONE 6 (5): e19539. https://doi.org/10.1371/journal.pone.0019539.

Shea, Charles H., Attila J. Kovacs, and Stefan Panzer. 2011. “The Coding and Inter-Manual Transfer of Movement Sequences.” Frontiers in Psychology 2 (April). https://doi.org/10.3389/fpsyg.2011.00052.

Trautmann, Eric M., Sergey D. Stavisky, Subhaneil Lahiri, Katherine C. Ames, Matthew T Kaufman, Daniel J. O’Shea, Saurabh Vyas, et al. 2019. “Accurate Estimation of Neural Population Dynamics without Spike Sorting.” Neuron, 2019, In press edition.

Trautmann, Eric M., Sergey D. Stavisky, Subhaneil Lahiri, Katherine C. Ames, Matthew T. Kaufman, Stephen I. Ryu, Surya Ganguli, and Krishna V. Shenoy. 2017. “Accurate Estimation of Neural Population Dynamics without Spike Sorting.” BioRxiv, December, 229252. https://doi.org/10.1101/229252.

Wesselink, Daan B, Fiona MZ van den Heiligenberg, Naveed Ejaz, Harriet Dempsey-Jones, Lucilla Cardinali, Aurelie Tarall-Jozwiak, Jörn Diedrichsen, and Tamar R Makin. 2019. “Obtaining and Maintaining Cortical Hand Representation as Evidenced from Acquired and Congenital Handlessness.” Edited by Eve Marder. ELife 8 (February): e37227. https://doi.org/10.7554/eLife.37227.

Wiestler, Tobias, Sheena Waters-Metenier, and Jörn Diedrichsen. 2014. “Effector-Independent Motor Sequence Representations Exist in Extrinsic and Intrinsic Reference Frames.” The Journal of Neuroscience: The Official Journal of the Society for Neuroscience 34 (14): 5054–64. https://doi.org/10.1523/JNEUROSCI.5363-13.2014.

Yokoi, Atsushi, Wenjun Bai, and Jörn Diedrichsen. 2016. “Restricted Transfer of Learning between Unimanual and Bimanual Finger Sequences.” Journal of Neurophysiology 117 (3): 1043–51. https://doi.org/10.1152/jn.00387.2016.

Yousry, T. A., U. D. Schmid, H. Alkadhi, D. Schmidt, A. Peraud, A. Buettner, and P. Winkler. 1997. “Localization of the Motor Hand Area to a Knob on the Precentral Gyrus. A New Landmark.” Brain 120 (1): 141–57. https://doi.org/10.1093/brain/120.1.141.

Zhang, Carey Y., Tyson Aflalo, Boris Revechkis, Emily R. Rosario, Debra Ouellette, Nader Pouratian, and Richard A. Andersen. 2017. “Partially Mixed Selectivity in Human Posterior Parietal Association Cortex.” Neuron 95 (3): 697-708.e4. https://doi.org/10.1016/j.neuron.2017.06.040.

Zilles, K., G. Schlaug, M. Matelli, G. Luppino, A. Schleicher, M. Qü, A. Dabringhaus, R. Seitz, and P. E. Roland. 1995. “Mapping of Human and Macaque Sensorimotor Areas by Integrating Architectonic, Transmitter Receptor, MRI and PET Data.” Journal of Anatomy 187 (Pt 3) (December): 515–37.

